# Acquisition of alveolar fate and differentiation competence by human fetal lung epithelial progenitor cells

**DOI:** 10.1101/2021.06.30.450501

**Authors:** Kyungtae Lim, Walfred Tang, Dawei Sun, Peng He, Sarah A. Teichmann, John C. Marioni, Kerstin B. Meyer, Emma L. Rawlins

**Affiliations:** Wellcome Trust/CRUK Gurdon Institute, Department of Physiology, Development and Neuroscience, Wellcome Trust/MRC Stem Cell Institute, University of Cambridge, Cambridge, CB2 1QN, UK; Wellcome Sanger Institute, Hinxton, Cambridge, CB10 1SA, UK; European Molecular Biology Laboratory, European Bioinformatics Institute (EMBL-EBI), Wellcome Genome Campus, Cambridge, UK

**Keywords:** human lung development, distal tip, organoids, alveolar differentiation, NKX2.1, stem cell, Wnt

## Abstract

Variation in lung alveolar development is strongly linked to disease susceptibility. However, the cellular and molecular mechanisms underlying alveolar development are difficult to study in humans. Using primary human fetal lungs we have characterized a tip progenitor cell population with alveolar fate potential. These data allowed us to benchmark a self-organising organoid system which captures key aspects of lung lineage commitment and can be efficiently differentiated to alveolar type 2 cell fate. Our data show that Wnt and FGF signalling, and the downstream transcription factors NKX2.1 and TFAP2C, promote human alveolar or airway fate respectively. Moreover, we have functionally validated cell-cell interactions in human lung alveolar patterning. We show that Wnt signalling from differentiating fibroblasts promotes alveolar type 2 cell identity, whereas myofibroblasts secrete the Wnt inhibitor, NOTUM, providing spatial patterning. Our organoid system recapitulates key aspects of human lung development allowing mechanistic experiments to determine the underpinning molecular regulation.

## INTRODUCTION

During human lung development the airway tree is formed by branching between ∼5 and 16 post conception weeks (pcw) in the pseudoglandular stage of development. At the canalicular stage, ∼16 to 26 pcw, the most distal epithelial tubes narrow, come into close proximity to capillaries and start to differentiate as alveolar epithelial cells^1,2^. Preterm infants born during the late canalicular stage have a rudimentary gas exchange surface and can survive if provided with specialised, neonatal intensive care. However, the molecular mechanisms underlying human alveolar development remain largely unknown.

During airway branching, the human tip epithelium is SOX9, SOX2 dual-positive and functions as a multipotent progenitor. Pseudoglandular tip epithelium has been cultured as self-renewing organoids which model the airway branching stage^3,4^. During the canalicular stage, as alveolar differentiation begins, tip progenitors become SOX9 single-positive and more cuboidal in shape^3^. We hypothesized that growth of tip organoids from canalicular stage lungs would provide an improved model for studying human alveolar differentiation.

Differentiation of human iPSCs to alveolar lineages suggests Wnt signalling is essential for alveolar fate^5^. NKX2.1 is also implicated in alveolar differentiation. In mouse lungs, Nkx2.1 is essential for alveolar differentiation and maintenance and binds to promoters of alveolar type 1 (AT1) and alveolar type 2 (AT2) cell-specific genes^6^. Heterozygous missense mutations in the *NKX2.1* homeodomain cause brain-lung-thyroid syndrome which includes disrupted surfactant gene expression and interstitial lung disease^7,8^. However, whether NKX2.1 simply promotes surfactant synthesis, or has additional roles in human lung alveolar differentiation is currently unknown.

We find that SOX9^+^ human lung tip progenitors acquire an AT2 gene expression signature by the canalicular stage of development. We derive, and characterise, a canalicular stage tip self-renewing organoid model. This has allowed us to determine upstream signals and downstream TFs which promote lung lineage commitment and investigate the spatial patterning of the developing alveolus.

## RESULTS

### Human fetal lung tip progenitor cells acquire alveolar lineage signatures *in vivo*

We investigated the distal lung tip epithelium of human fetal lungs at the pseudoglandular and canalicular stages. Consistent with our previous report^3^, pseudoglandular distal tips were columnar and marked by SOX9 and SOX2. The canalicular stage distal tip epithelium contained more cuboidal SOX2^-^ SOX9^+^ cells which co-expressed the AT2 cell markers, SFTPC and HTII-280 (Fig. 1A,B). We identified surface markers to distinguish between pseudoglandular and canalicular stage tips. CD44 marks tip epithelial cells across all stages of lung development tested. Whereas CD36 is expressed specifically in the canalicular stage tip (∼16-21 pcw) where it is co-expressed with CD44, *SFTPC* and SOX9. A lower level of both CD44 and CD36 extends into the SOX9^-^/PDPN^+^ tip-adjacent cells and CD44 extends further proximally into the differentiating stalk region (Figs. 1C-F; Extended Data Fig. 1A,B).

**Fig. 1.**
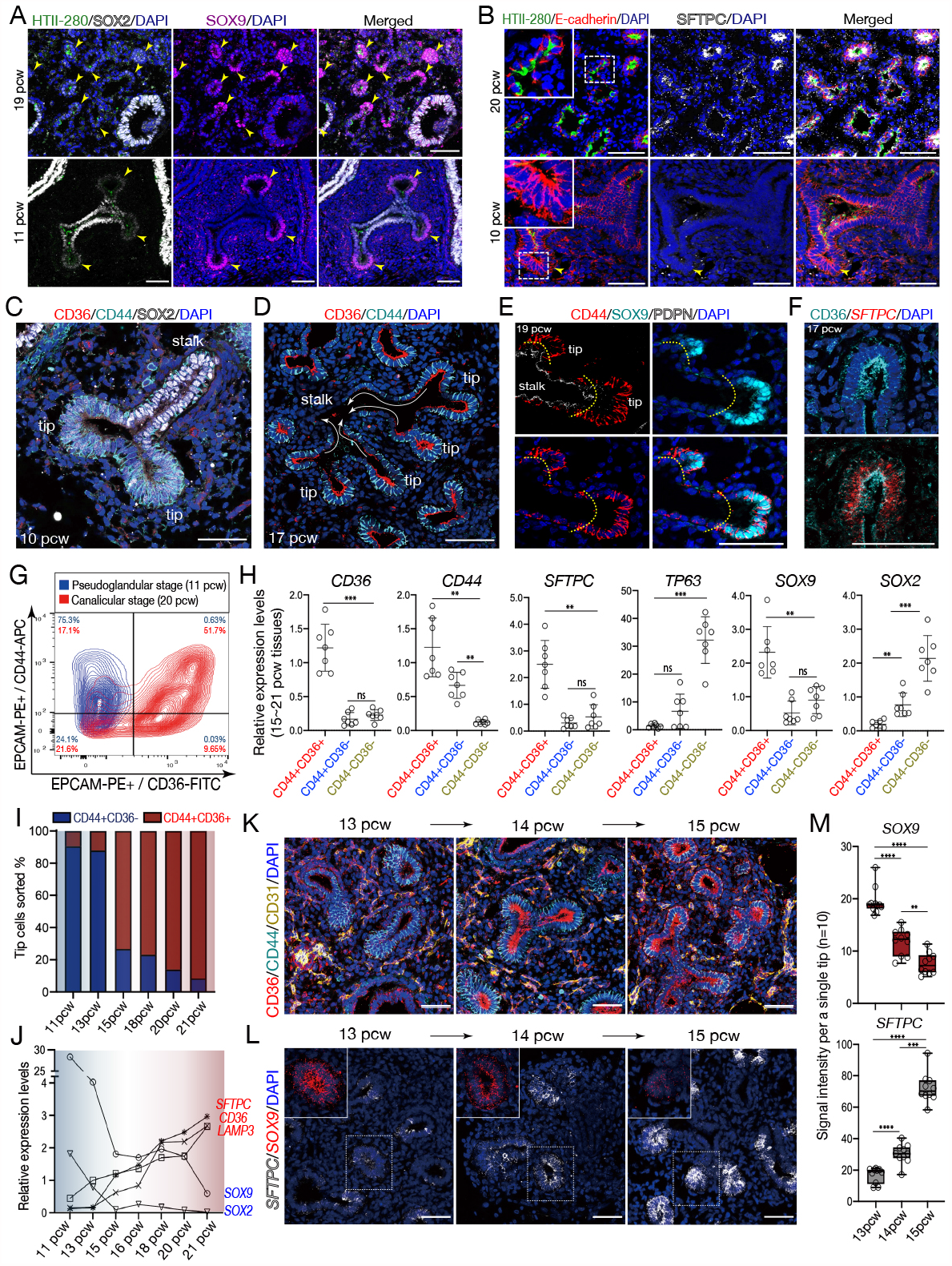
Human fetal lung tip progenitor cells acquire alveolar features during normal development. (A and B) Human fetal lung at pseudoglandular and canalicular stages; 11, 19 pcw (A) and 10, 20 pcw (B). Tip epithelium (arrowheads) is marked by E-cadherin, SFTPC, HTII-280, and SOX9. SOX2, airway epithelium (A). (C-F) Surface antigens, CD44 and CD36, mark tip epithelium at pseudoglandular and canalicular stages. Lungs at 10 (C), 17 (D, F), and 19 pcw (E) were stained with CD36 and CD44 and/or SOX2 and SOX9 antibodies. Arrows (D) show patterning from distal to proximal regions. Yellow dashed lines (E) indicate separation of SOX9^+^ tip regions from the PDPN^+^ stalk. The *SFTPC* transcript was visualised by *in situ* HCR (F) following immunostaining for CD36. (G) Flow cytometry of the human lung tip epithelial population at the pseudoglandular and canalicular stages, 11 (*blue*) and 20 pcw (*red*). (H) qRT-PCR of the freshly purified lung epithelial cells from canalicular stage lungs sorted. Data normalized to fresh EPCAM+ cells from 20 pcw distal tissues; mean ± SD, *n* = 7 (15∼21 pcw). Significance evaluated by 1-way ANOVA with Tukey multiple comparison post-test; ns: not significant, \**P*<0.05, \*\**P*<0.01, \*\*\**P*<0.001. (I) Proportion of the freshly purified tip epithelium as CD44^+^CD36^-^ or CD44^+^CD36^+^ at 11, 13, 15, 18, 20 and 21 pcw; *n* = 1 each time. (J) Relative mRNA levels of the tip progenitor markers, *SOX9* and *SOX2*, and type 2 alveolar lineage markers, *SFTPC, CD36* and *LAMP3*, in CD44^+^CD36^-^ tip epithelial population at 11 and 13 pcw, and in CD44^+^CD36^+^ tip epithelial population at 15, 16, 18, 20 and 21 pcw, by qRT-PCR. Data was normalized to fresh EPCAM^+^ cells from 20 pcw tip tissues; *n* = 1 at each stage. (K and L) Human fetal lung tissues during the transition from 13 to 15 pcw were stained using antibodies against CD36, CD44, and CD31 (K), or for *SFTPC* and *SOX9* mRNA (L). Three 13 pcw, two 14 pcw, and two 15 pcw samples. (M) Signal intensity of *SOX9* and *SFTPC* transcripts in Fig. 1L. Ten tip regions analysed per stage and the intensity represented as mean ± SD. Significance evaluated by 1-way ANOVA with Tukey multiple comparison post-test; ns: not significant, \**P*<0.05, \*\**P*<0.01, \*\*\**P*<0.001, \*\*\*\**P*<0.001. DAPI, nuclei. Scale bar, 50 μm.

Distal lung regions were dissected to enrich for the tip and EPCAM^+^ cells were sorted for CD44 and/or CD36. At the pseudoglandular stage (11 pcw), 75% of sorted cells were CD44^+^ and CD44^+^CD36^+^ cells were rare (Fig. 1G; *blue*). By contrast, at the canalicular stage (20 pcw) 53% of sorted cells were CD44^+^CD36^+^ and only 17% were single CD44^+^ (Fig. 1G; *red*). qRT-PCR showed that the 17-20 pcw CD44^+^CD36^+^ cells robustly expressed *CD36, CD44, SFTPC* and *SOX9*, but extremely low levels of the airway markers, *TP63* and *SOX2*, consistent with the immunostaining (Fig. 1H). In contrast, single CD44^+^ cells showed a higher level of *SOX2*, but much lower levels of *SFTPC* and *SOX9*, suggesting that they are derived from the CD44^+^SOX9^-^ stalk region (Fig. 1H; Extended Data Fig. 1B). Finally, the CD44^-^CD36^-^ cells had higher levels of TP63 and *SOX2*, but low *SFTPC* and *SOX9*, indicating they are derived from more proximal airway-lineage cells (Fig. 1H). Therefore, dual expression of CD44 and CD36 marks the tip epithelial population in the canalicular stage lung.

### Gradual acquisition of tip alveolar lineage signature during human lung development

Flow cytometric analysis showed that the expression of CD36 was robustly acquired between 13 and 15 pcw, prior to the canalicular stage (Fig. 1I). Similarly, in the CD36^+^ cells, the mRNA levels of *SFTPC, CD36*, and *LAMP3* began to increase from 13 pcw onward (Fig. 1J). We confirmed that CD36 was detectable in the tip epithelium at exactly 14 pcw, moreover the intensity of *SFTPC* transcripts increased while *SOX9* gradually lowered during this transition period (Fig. 1K-M). These results show that the acquisition of AT2 lineage signatures occurs gradually in the tip epithelium prior to the canalicular stage.

### Organoids derived from distal tip epithelium at the canalicular stage exhibit alveolar lineage signatures

To determine the fate potential of the canalicular stage (17-20 pcw) distal tip, CD44^+^CD36^+^ cells were cultured for 3 weeks (Fig. 2A). Two morphologically distinct organoids formed: cystic and folded (Fig. 2B). All the folded organoids consisted of cuboidal cells and expressed both progenitor and AT2 markers, including an *SFTPC*-eGFP reporter (Fig. 2C). By contrast, the cystic organoids had a columnar cell shape and expressed tip progenitor markers, but not AT2 markers, resembling the pseudoglandular tip epithelium (Fig. 2B-E; Extended Data Fig. 2A,B). For simplicity, we refer to the folded and cystic organoids isolated from 17-21 pcw lungs as lineage positive (Lin^POS^) and negative (Lin^NEG^).

**Fig. 2.**
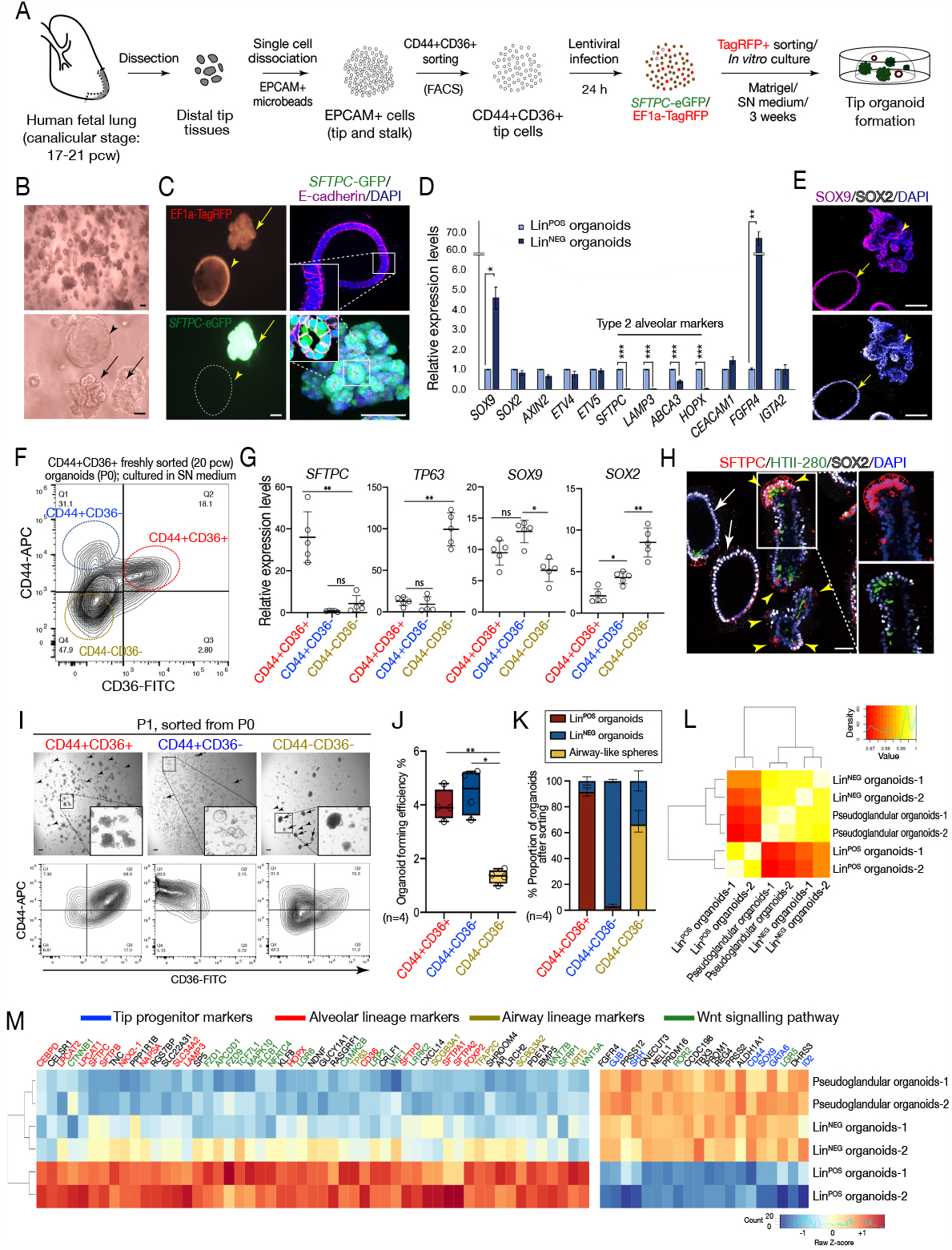
CD36, CD44 dual-positive tip cells self-renew and undergo lineage commitment *in vitro* to form canalicular stage lung organoids. (A) Isolation and viral infection of CD44^+^CD36^+^ tip epithelial cells from human fetal lungs at the canalicular stage and *in vitro* culture in self-renewing (SN) medium. (B and C) Gross morphology (B) of the cultured epithelial tip organoids. Detailed morphology (C) E-cadherin (magenta); *SFTPC-*GFP and *TagRFP*. Arrows and arrowhead indicate folded Lin^POS^ organoids and cystic Lin^NEG^ organoids, respectively. Scale bars, 100 μm. (G) Gene expression profile of the Lin^POS^ and Lin^NEG^ organoids. Data are quantified by qRT-PCR; mean ± SD of 4 biological replicates. Significance evaluated by unpaired student *t*-test; \**P*<0.05, \*\**P*<0.01, \*\*\**P*<0.001. (E) Immunofluorescence analysis of the Lin^POS^ (arrowheads) and Lin^NEG^ organoids (arrow) at passage 1 cultured in the SN medium, showing co-expression of SOX9 and SOX2. DAPI, nuclei. Scale bar, 50 μm. (F and G) Canalicular stage lung tip organoids sorted into 3 populations at passage zero by FACS using antibodies against CD36 and CD44 (F). The sorted P0 cell populations were analysed by qRT-PCR (G). Data was normalized to total EPCAM^+^ cells freshly sorted from 20 pcw tissues; mean ± SD (*n* = 5). Significance was evaluated by 1-way ANOVA with Tukey multiple comparison post-test; \**P*<0.05, \*\**P*<0.01. (H) Tip organoids at passage zero cultured in self-renewal medium stained with SFTPC, HTII-280 and SOX2 antibodies. Arrowheads indicate the tip-like, SFTPC^+^ subpopulation in the Lin^POS^ organoids. Arrows indicate the Lin^NEG^ organoids. (I-K) Passage 1 organoids were grown from the sorted CD44^+^CD36^+^, CD44^+^CD36^-^ or CD44^-^ CD36^-^ populations at Passage 0 and reanalysed for CD44 and CD36 at the end of Passage 1 (I; Extended Data Fig. 2L). Arrows and arrowheads indicate the Lin^POS^ and Lin^NEG^ organoids. The organoid forming efficiency (J) and the proportion (K) of the organoids of each morphological sub-type at passage 1 was measured at 3 weeks after plating. Data was represented as mean ± SD of 4 biological replicates. Scale bar, 100 μm. (L) Hierarchical clustering analysis of bulk-RNA seq data using pseudoglandular, Lin^POS^ and Lin^NEG^ organoids. (M) Heatmap analysis of selected genes highly enriched in pseudoglandular, Lin^NEG^ and Lin^POS^ organoids.

We tested whether any organoids (from the mixed Lin^POS^ and Lin^NEG^ population) retained CD44 and CD36 expression after 3 weeks culture (Fig. 2F). The CD44^+^CD36^+^ cells showed the highest level of *SFTPC* with a moderate level of *SOX9*, but very low levels of *SOX2* and *TP63*. They were located at the tips of the Lin^POS^ organoids where they expressed SFTPC, HTII-280, SOX9, CD44 and KI67 (Fig. 2G,H; Extended Data Fig. 2C-F). The CD44^-^CD36^-^ cells had the highest levels of *TP63* and *SOX2*. They corresponded to the inner parts of the Lin^POS^ organoids where scattered TP63^+^ cells were found (Fig. 2G,H; Extended Data Fig. 2D). By contrast, the CD44^+^CD36^-^ cells had the highest level of *SOX9* and a moderate level of *SOX2*, but no lineage markers, and corresponded to the Lin^NEG^ organoids which had uniform CD44, SOX2 and SOX9 (Fig. 2G,H; Extended Data Fig. 2E). These data suggested that the CD44^+^CD36^+^ canalicular stage tip cells originally plated had self-renewed (at the tips) and differentiated towards airway lineages (in the centre) to form the Lin^POS^ organoids. Moreover, that a fraction of the CD44^+^CD36^+^ tip cells had also grown into Lin^NEG^ organoids, resembling the pseudoglandular stage tips. To test this hypothesis, we infected freshly-isolated CD44^+^CD36^+^ epithelial cells with *SFTPC*-eGFP^+^ lentivirus and sorted for eGFP. We observed that sorted *SFTPC*-eGFP^+^ cells formed both Lin^POS^ and Lin^NEG^ organoids (Extended Data Fig. 2G-I). These data suggest that the emergence of Lin^NEG^ organoids, which are similar to pseudoglandular stage tip organoids, is due to dedifferentiation of the epigenetically unstable canalicular tip epithelium. However, we cannot exclude the possibility that the canalicular tip epithelium contains a mixture of cell states.

We tested if the CD44^+^CD36^+^ cells continued to self-renew upon passaging. P0 organoids (mixed population of Lin^POS^ and Lin^NEG^) were sorted as CD44^+^CD36^+^, CD44^+^CD36^-^ and CD44^-^CD36^-^ and cultured separately in the self-renewal medium. Only the CD44^+^CD36^+^ cells were able to generate a large proportion of Lin^POS^ organoids with folded structure and progenitor/AT2 gene signature (Figs. 2I-K; Extended Data Fig. 2J-L). In contrast, the CD44^+^CD36^-^ cells, derived from Lin^NEG^ organoids, produced Lin^NEG^ organoids. The CD44^-^ CD36^-^ cells, derived from the centre of the Lin^POS^ organoids, largely formed airway-fated spheres expressing a significantly higher level of *TP63/*TP63 (Fig. 2I-K; Extended Data Fig. 2J-L). These data conclusively demonstrate that the CD44^+^CD36^+^ cells are the major tip progenitor subpopulation *in vitro* and can maintain the Lin^POS^ organoids. We have therefore captured the canalicular stage lung tip epithelial population which co-expresses SOX9 and AT2 markers in the Lin^POS^ organoids.

We performed RNAseq to compare the transcriptome of passaged pseudoglandular stage tip organoids derived from 8-9 pcw^3^ with passaged canalicular stage Lin^NEG^ and Lin^POS^ organoids (Extended Data Fig. 3A). Hierarchical clustering and principal component analysis showed that the Lin^NEG^ organoids were very similar to the pseudoglandular organoids, but distinct from the Lin^POS^ organoids (Fig. 2L; Extended Data Fig. 3B). We identified >280 differentially expressed genes between the Lin^POS^ and Lin^NEG^ organoids (Supplementary Table 1; Extended Data Fig. 3C; log_2_FC>4, *P*<0.05). Similar to the pseudoglandular organoids, the Lin^NEG^ organoids were highly associated with Gene Ontology (GO) terms related to ion transport and branching morphogenesis, confirming that they resemble the pseudoglandular stage distal tips. Whereas the Lin^POS^ organoids had significant GO terms for respiratory gaseous exchange and lung alveolus development, as well as canonical Wnt pathway signalling (Extended Data Fig. 3D-G). Moreover, the Lin^POS^ organoids were enriched for AT2 markers *SFTPC, SLC34A2, NAPSA, LPCAT1, FOXP2*, and *CEBPD*, Wnt signalling-related genes *CTNNB1, TCF7L1, WNT7B, WIF1, LRRK2* and low levels of airway genes *TP63, SCGB3A1, SCGB3A2* (Fig. 2M; Supplementary Table 1). These data confirm that the passaged Lin^POS^ organoids recapitulate key molecular characteristics of the canalicular stage lung tip epithelial progenitors.

### Coordinated control of the canalicular stage tip epithelial cell fate by Wnt and FGF signalling

To determine which signalling cues direct differentiation of the canalicular tips, tip epithelium was isolated from the distal lung and directly exposed to pairwise signal combinations (Fig. 3A; Extended Data Fig. 4A). The cells did not grow in the absence of SMADi (Noggin and SB431542) (Fig. 3B; Extended Data Fig. 4B). However, we observed that two distinct populations of organoids could be obtained by combining SMADi with CHIR (CHIR99021, a Wnt agonist), or with FGFs (FGF7 and FGF10) (Fig. 3B). Organoids grown in SMADi/CHIR had a thin epithelium with a hollow lumen. They could not be passaged and expressed the highest level of *SFTPC*, greater than the Lin^POS^ organoids. In contrast, organoids grown in SMADi/FGF formed spheres with a small lumen, a relatively thick, proliferative epithelial layer and expressed the highest level of TP63 (Fig. 3B-D). In both conditions *SOX9* levels were lower than the Lin^POS^ and Lin^NEG^ organoids (Fig. 3C). These data indicate that Wnt and FGF signalling promote the lineage determination of the canalicular stage tip epithelium to alveolar or airway lineages, in agreement with previously published data^5,9^. Moreover, when SMADi/CHIR/FGFs were combined (equivalent to our self-renewing medium) organoids displayed a mixture of alveolar and airway characteristics, as in the Lin^POS^ organoids (Fig. 3D).

**Fig. 3.**
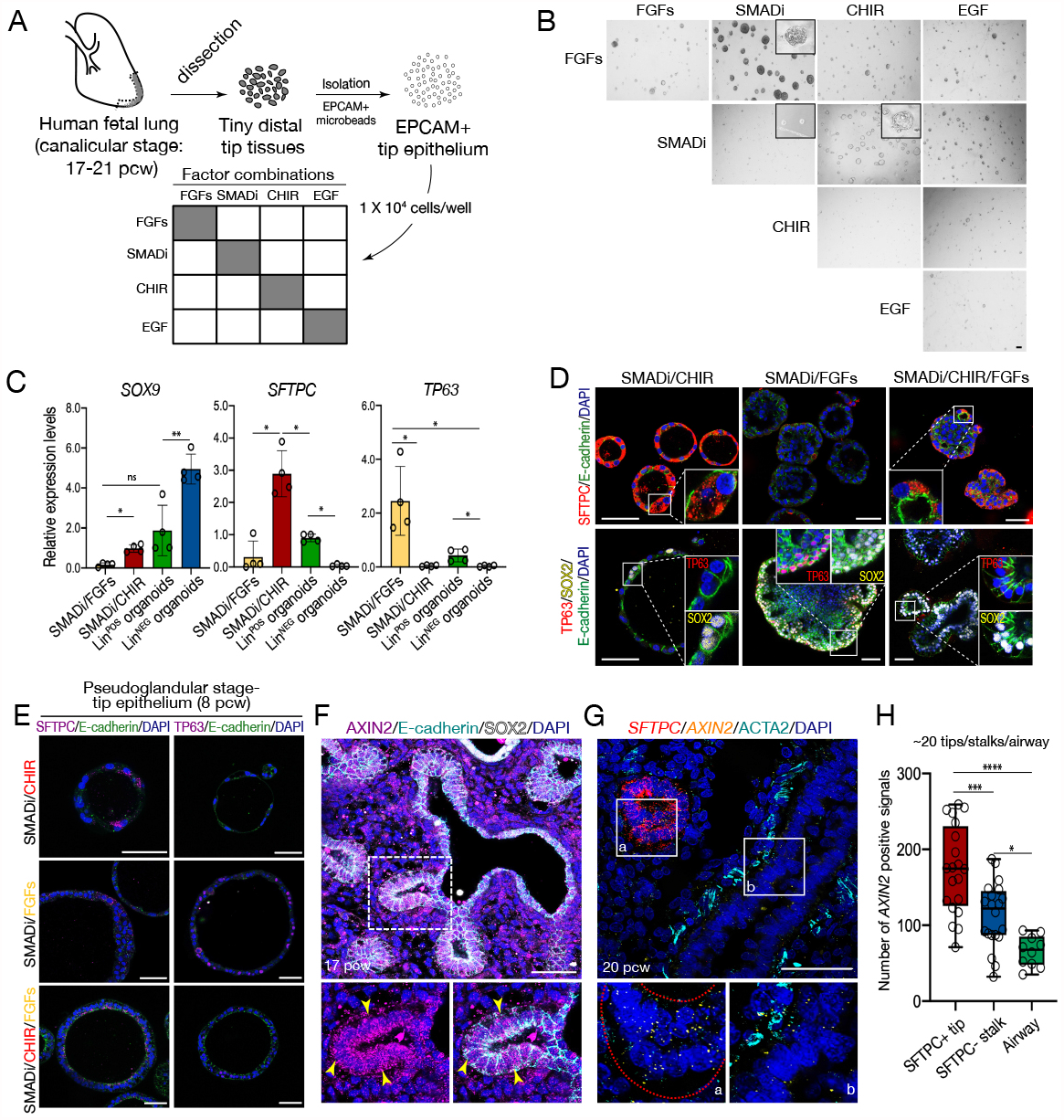
Wnt and FGF signalling coordinate human tip cell maintenance and differentiation *in vitro*. (A) Diagram showing *in vitro* culture of the tip epithelial cells for 2 weeks in single or pairwise combinations of signalling cues: FGFs (FGF7, FGF10), SMADi (Noggin, SB431542), CHIR (CHIR99021) and EGF. (B) Morphology of the tip organoids cultured in different culture conditions for 2 weeks. Representative image from 1 biological replicate is shown; n=4 biological replicates in total. Scale bar, 200 μm. (C) Relative mRNA levels of *SOX9, SFTPC* and TP63 measured by qRT-PCR. Normalized to a Lin^POS^ organoid line; mean ± SD of 4 independent biological replicates. Significance was evaluated by 1-way ANOVA with Tukey multiple comparison post-test; ns: not significant, \**P*<0.05 and \*\**P*<0.01. (D) Immunofluorescence analysis of the tip organoids at passage zero cultured in the different culture conditions. Antibodies against E-cadherin, SFTPC, TP63 and SOX2 were used. Scale bar, 20 μm. (E) Immunofluorescence analysis of the pseudoglandular organoids derived from an 8 pcw pseudoglandular stage lung in the different conditions at passage 0. Antibodies against SFTPC, TP63, E-cadherin and SOX2 were used. Scale bar, 50 μm. (F-H) Frozen sections of human fetal lung tissues at 17 pcw (F) and 20 pcw (G) were immunostained for AXIN2, E-cadherin and SOX2 (F), or ACTA2 followed by *in situ* HCR for *SFTPC* and *AXIN2* (G). Arrowheads (F) indicate AXIN2^+^ tip epithelial cells. Red dashed line in the inset (G) indicates SFTPC^+^ tip epithelial cells. *AXIN2* signals were counted from 20 areas of tips and stalks and 10 areas of airway across 3 independent lung tissues at 18-20 pcw (H). Significance was evaluated by 1-way ANOVA with Tukey multiple comparison post-test; ns: not significant, \**P*<0.05, \*\**P*<0.01, \*\*\**P*<0.001 and \*\*\*\**P*<0.0001. Scale bar, 50 μm. DAPI indicates nuclei.

We demonstrated that the canalicular stage tips are highly plastic and can switch readily between alveolar and airway differentiation by altering the medium and observing rapid organoid morphology and gene expression changes (Extended Data Fig. 4C-E). Moreover, FGF7 alone, without FGF10, was sufficient to promote airway fate (Extended Data Fig. 4F,G). Freshly-isolated pseudoglandular stage (8 pcw) distal tip epithelial cells did not show similar levels of differentiation when grown in the same conditions (Fig. 4H). These data indicate that the canalicular stage tip progenitors are both highly plastic and in a differentiation-ready state compared to the pseudoglandular tip.

**Fig. 4.**
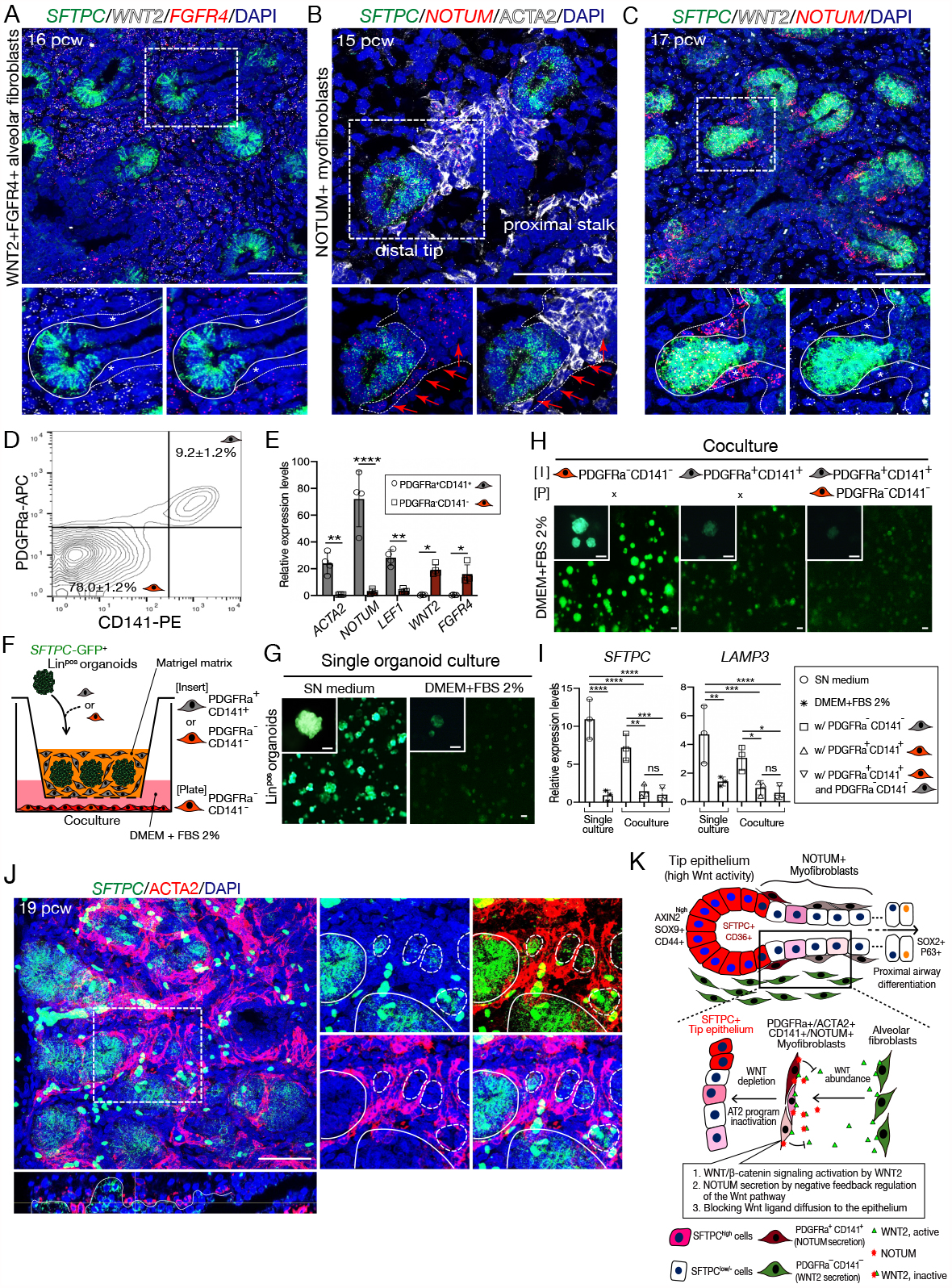
Spatial patterning of the differentiating alveolar epithelium by NOTUM+ myofibroblasts. (A-C) Frozen sections of 15-17 pcw human fetal lung stained by *in situ* HCR and/or antibodies. A. 16 pcw, *SFTPC, WNT2*, and *FGFR4* probes. B. 15 pcw, *SFTPC* and *NOTUM* probes, ACTA2 antibody. C. 17 pcw *SFTPC, WNT2*, and *NOTUM* probes. Arrows (B) and asterisks (A,C) represent ACTA2^+^*NOTUM*^+^ myofibroblasts in the tissues. Lines and dashed lines indicate the boundaries of epithelial cells and myofibroblasts, respectively. Scale bar, 50 μm. (D) Isolation of PDGFRA^+^CD141^+^ myofibroblasts and PDGFRA^-^CD141^-^ alveolar fibroblasts from human fetal lung tissues at 17-21 pcw using a combination of PDGFRa-APC and CD141-PE antibodies. (E) qRT-PCR of PDGFRa^+^CD141^+^ myofibroblasts and PDGFRa^-^CD141^-^ alveolar fibroblasts freshly isolated from 17 to 21 pcw human lung tissues. Data was normalized to the total isolated fibroblast population; mean ± SD of biological 4 replicates. Significance was evaluated by unpaired student *t*-test; \**P*<0.05, \*\**P*<0.01, \*\*\**P*<0.001. (F) Diagram illustrating *in vitro* coculture of the isolated PDGFRa^+^CD141^+^ myofibroblasts and PDGFRa^-^CD141^-^ alveolar fibroblast with Lin^POS^ tip organoids expressing *SFTPC-GFP*. (G and H) *SFTPC-GFP* signal of Lin^POS^ tip organoids. G. Cultured alone in self-renewing (SN) or DMEM + 2% FBS medium. H. Cocultured with freshly isolated fibroblast sub-populations. I, insert; P, plate. Scale bar, 200 μm. (I) qRT-PCR for *SFTPC* and *LAMP3*, 2 weeks after *in vitro* culture. Mean ± SD of 3 biological replicates. Significance was evaluated by one-way ANOVA; ns: not significant, \**P*<0.05, \*\**P*<0.01, \*\*\**P*<0.001. (J) Thick sections of human fetal lung immunostained with ACTA2 followed by *in situ* HCR for *SFTPC*. Lines and dashed lines in the inset indicate *SFTPC*^+^ epithelial cell populations located in tip and stalk regions, respectively. Thickness: 42 μm. See also Supplementary Video 1. (K) Summary diagram showing the spatial regulation of Wnt signalling mediated by ACTA2^+^PDGFRa^+^CD141^+^ myofibroblasts in the distal regions of human lung tissues during the canalicular stage. DAPI indicates nuclei. Scale bar, 50 μm.

We reasoned that Wnt and FGF signalling likely control lineage determination of the tip epithelium *in vivo*. In canalicular stage tissue, we found that the *SFTPC*^+^ tips expressed higher levels of the Wnt targets *AXIN2* and *WIF1*, compared with stalk and airway epithelium (Figs. 3F-H; Extended Data Figs. 4H,I; 5A-D). We observed that *WNT2* is co-expressed with *FGFR4* in fibroblasts throughout the canalicular stage (Fig. 4A); putative alveolar fibroblasts^10^. This led us to question how Wnt-responsive *SFTPC* could be precisely restricted to the tip epithelium in the presence of widespread *WNT2*. A secreted Wnt inhibitor, *NOTUM*^11^, is expressed in both the distal tip epithelium and the myofibroblast/smooth muscle population which surrounds the differentiating epithelial stalk cells (Fig. 4B,C; Extended Data Fig. 5B,C). The *NOTUM*^*+*^ myofibroblasts co-express the Wnt targets *LEF1* and *AXIN2*, suggesting that they may also be responding to Wnt (Extended Data Fig. 5C,D). We hypothesised that in response to the WNT2 signal, the stalk myofibroblasts locally secrete NOTUM, preventing the stalk epithelium from experiencing a high level of Wnt activity, allowing them to turn off *SFTPC* and exit the tip fate. We identified surface antigens for the specific isolation of WNT2^+^ fibroblasts or NOTUM^+^ myofibroblasts (Fig. 4D; Extended Data Fig. 5E,F). Isolated PDGFRA^+^CD141^+^ myofibroblasts express high levels of *ACTA2, NOTUM* and *LEF1*. Whereas PDGFRA^-^CD141^-^ fibroblasts express high levels of *WNT2* and *FGFR4* (Fig. 4E). This specific gene expression is maintained if the cell types are cultured individually for 14 days. When freshly isolated fibroblasts and myofibroblasts were co-cultured, the levels of *LEF1* and *NOTUM* expression increased in the myofibroblasts, suggesting that they are indeed responding to WNT2 from the fibroblasts (Extended Data Fig. 5G).

We therefore asked whether co-culture with the PDGFRA^-^CD141^-^ fibroblasts, or PDGFRA^+^CD141^+^ myofibroblasts, could affect *SFTPC* expression in the Lin^POS^ organoids (Fig. 4F). Lin^POS^ organoids robustly express the *SFTPC*-GFP reporter when cultured in the self-renewing medium, but not in 2% FBS (Fig. 4G). However, co-culture with PDGFRA^-^ CD141^-^ fibroblasts is sufficient to substitute for the self-renewing medium and maintain *SFTPC*-GFP and endogenous *SFTPC* and *LAMP3* (Fig. 4H,I). By contrast, co-culture of the Lin^POS^ organoids with PDGFRA^+^CD141^+^ myofibroblasts, or both PDGFRA^+^CD141^+^ myofibroblasts and PDGFRA^-^CD141^-^ fibroblasts, did not support expression of AT2 genes. This leads us to propose that *in vivo* WNT2-expressing alveolar fibroblasts promote *SFTPC* expression in the distal tip. Moreover, that differentiating stalk cells are protected from the Wnt signal by the NOTUM-secreting myofibroblasts allowing them to turn off *SFTPC* and enter a differentiation programme (Fig. 4K).

Differentiating AT2 cells are also *SFTPC*^+^. We observed that the differentiating *SFTPC*^+^ cells were never directly over-lain by the *NOTUM*^*+*^ myoepithelial cells (Fig. 4J). This further supports the concept that the differentiating alveolar epithelium is patterned by signals from the myofibroblasts (Fig. 4K).

### NKX2.1 is a major driving force for alveolar differentiation in the tip epithelial organoids

To identify putative transcription factors for cell differentiation we analysed chromatin accessibility of the pseudoglandular and Lin^POS^ organoids by bulk-ATAC seq. There were ∼2-fold more differentially open chromatin regions in the Lin^POS^ than pseudoglandular organoids, consistent with the increased cell type complexity of the Lin^POS^ organoids (Extended Data Fig. 6A; Supplementary Table 2). The genomic distribution of the differentially opened chromatin was similar in both organoid types (Extended Data Fig. 6B). GO analysis of the genes nearest to differentially open chromatin was consistent with the RNA-seq data (Fig. 2; Extended Data Fig. 6C-E). However, a much higher proportion of lung development-associated genes had open chromatin at the promoter regions in the Lin^POS^ organoids (Fig. 5A). For example, the promoter regions of lung differentiation genes *SFTPC, TP63, FOXP2* and *CD36* and Wnt signalling genes, *AXIN2, CTNNB1, DVL3, LRRK2* and *WIF1*, were more accessible in the Lin^POS^ than pseudoglandular organoids (Fig. 5B; Extended Data Fig. 6D,E). These data strongly suggest that the chromatin accessibility of the Lin^POS^ organoids is more favourable for lineage differentiation than the pseudoglandular organoids.

**Fig. 5.**
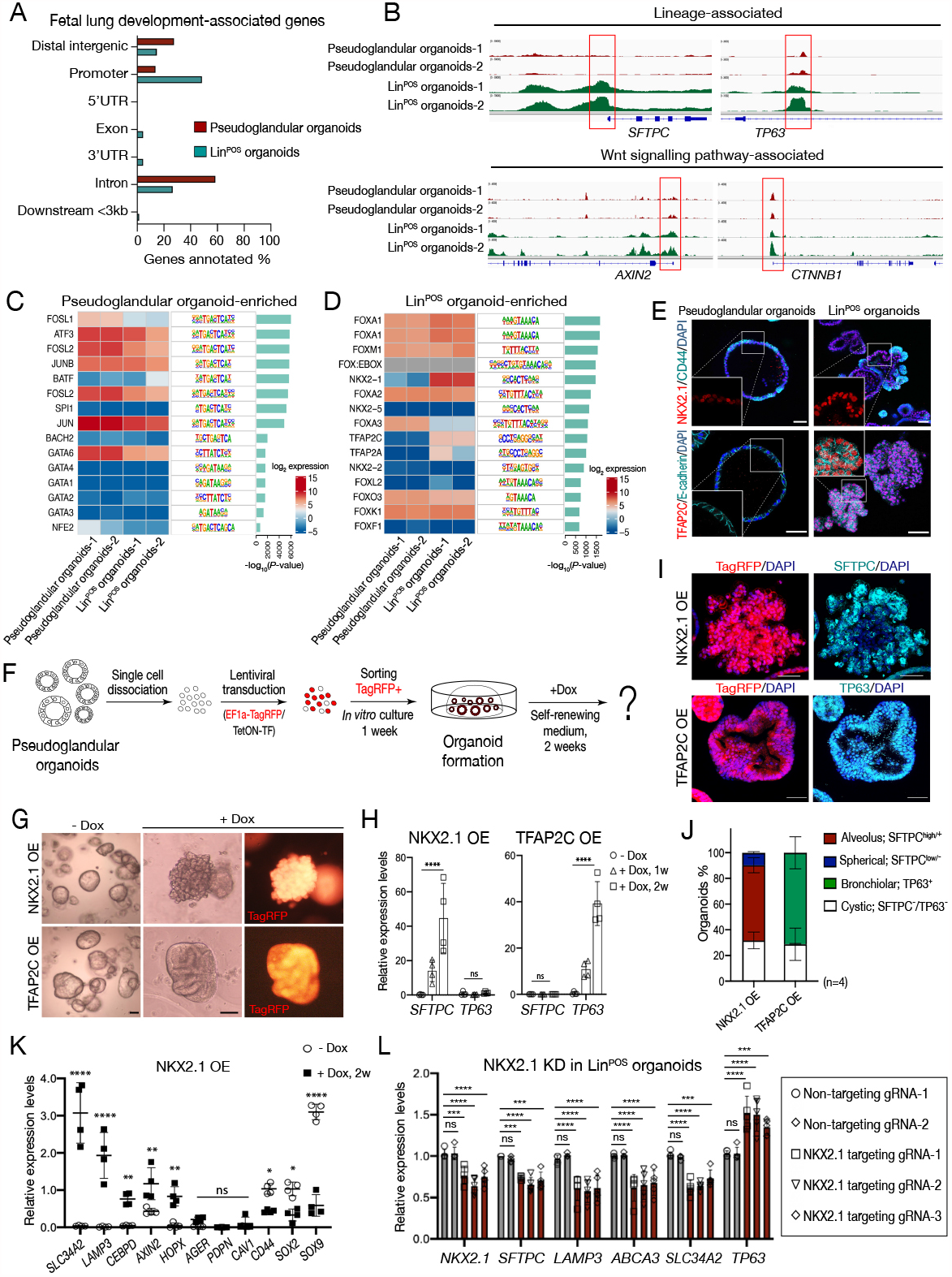
Identification of key transcription factors controlling airway and alveolar lineage differentiation using the organoid system. (A) Genomic distribution of differentially accessible chromatin regions associated with human fetal lung development between the pseudoglandular and Lin^POS^ organoids. (B) Representative ATAC-seq tracks visualized in Integrative Genomics Viewer (IGV) at *SFTPC, TP63, AXIN2* and *CTNNB1. Red* box indicates the promoter. (C and D) HOMER motif analysis coupled with RNA seq data. The top 15 most highly enriched motifs and TF gene expression level (heat map) are shown for the pseudoglandular (C) and Lin^POS^ organoids (D). (E) Pseudoglandular and Lin^POS^ organoids stained with antibodies against NKX2.1, TFAP2C, CD44 and E-cadherin. (F) Diagram showing doxycycline-inducible overexpression of NKX2.1 and/or TFAP2C in the pseudoglandular organoids. Constitutively expressed TagRFP was used for sorting transduced cells. (G) Morphology of the pseudoglandular organoids overexpressing NKX2.1 or TFAP2C for 2 weeks. Scale bar, 100 μm. (H) Relative mRNA levels of *SFTPC* and *TP63* were measured by qRT-PCR in NKX2.1- or TFAP2C-OE pseudoglandular organoids. Data was normalized to EPCAM^+^ cells freshly isolated from 20 pcw tip tissues; mean ± SD of four biological replicates. Significance was evaluated by 1-way ANOVA with Tukey multiple comparison post-test; ns: not significant, \*\*\*\**P*<0001. (I and J) SFTPC and TP63 antibody staining of the pseudoglandular organoids overexpressing NKX2.1 or TFAP2C for 2 weeks (I). The proportion of the organoids positively stained in (J) was measured based on morphology and signal intensity. N= 4 biological replicates. (K) qRT-PCR of the pseudoglandular organoids overexpressing NKX2.1 for 2 weeks. Data was normalized to EPCAM^+^ cells freshly isolated from 20 pcw tip tissues; mean ± SD of four biological replicates. Significance was evaluated by 1-way ANOVA with Tukey multiple comparison post-test; ns: not significant, \**P*<0.05, \*\**P*<0.01, \*\*\**P*<0.001 and \*\*\*\**P*<0.0001. (L) Knock-down (KD) of endogenous *NKX2.1* by CRISPR-dCas9-KRAB system. Data was normalized to EPCAM^+^ cells freshly isolated from 20 pcw tip tissues; mean ± SD of 5 biological replicates. Significance was evaluated by 1-way ANOVA with Tukey multiple comparison post-test; ns: not significant, \**P*<0.05, \*\**P*<0.01, \*\*\**P*<0.001, \*\*\*\**P*<0.0001. DAPI indicates nuclei. Scale bar, 50 μm.

To predict which transcription factors (TFs) control cell fate specification, we performed TF motif analysis in the differential ATAC seq peaks and compared this with our RNA seq data. The motifs for FOSL1 and GATA6 binding were differentially open, and *FOSL1* and *GATA6* highly expressed, in the pseudoglandular organoids (Fig. 5C). In the Lin^POS^ organoids, NKX2.1 and TFAP2C motifs were accessible, and these factors were highly expressed (Fig. 5D). Immunostaining confirmed that NKX2.1 is more strongly expressed in the Lin^POS^ than pseudoglandular organoids. Moreover, TFAP2C was absent in the pseudoglandular organoids, but ubiquitous in the Lin^POS^ organoids (Fig. 5E). *In vivo, NKX2-1* transcripts were most highly expressed in the alveolar regions of the lungs, whereas *TFAP2C* was expressed in the airway epithelium (Extended Data Fig. 7A,B).

We performed overexpression (OE) of NKX2.1 and TFAP2C in the pseudoglandular organoids to test if either factor was sufficient to induce differentiation to the alveolar or airway lineages (Fig. 5F). NKX2.1 OE resulted in ∼60% of the pseudoglandular organoids acquiring an alveolar-like structure with high levels of SFTPC expression (Fig. 5G-J; Extended Data Fig. 7C). NKX2.1 also significantly upregulated other AT2 lineage markers including *SCL34A2, LAMP3, CEBPD, HOPX* and *AXIN2*, but downregulated the tip markers *SOX9, SOX2* and *CD44* (Fig. 5K). In contrast, TFAP2C OE caused around 70% of the organoids to form bronchiolar-like structures and significantly increased basal cell markers including TP63/*P63, KRT5* and *NGFR*, (Fig. 5G-J; Extended Data Fig. 7C,D). Therefore, NKX2.1 and TFAP2C function as key regulators of differentiation toward AT2 and basal cell lineages respectively.

The Lin^POS^ organoid cells co-express high levels of NKX2-1 and TFAP2C (Fig. 5E) yet are comprised of distinct SOX9/SFTPC^+^ tip and TP63^Lo^ central regions (Fig. 2). We analysed the relationship between NKX2.1 and TFAP2C by overexpressing them together in the pseudoglandular organoids. The NKX2.1/TFAP2C OE organoids were highly folded, similar to NKX2-1 OE. Furthermore, TP63 was barely detectable, but SFTPC was markedly induced (Extended Data Fig. 7E). When Lin^POS^ organoids were cultured in SMADi/CHIR/FGF7, *NKX2.1* and *SFTPC* were high, but *TP63* and *TFAP2C* were low. Whereas in SMADi/FGF7 (without the Wnt agonist), NKX2.1 deceased ∼2-fold, *SFTPC* turned off and *TP63* and *TFAP2C* were robustly expressed (Extended Data Fig. 7F). These data clearly demonstrate that high NKX2.1 expression, in combination with Wnt signalling, suppresses the airway lineage program, also explaining why TP63 expression is low in the Lin^POS^ organoids although TFAP2C is expressed (Extended Data Fig. 7F). Further support for the importance of NKX2.1 in promoting alveolar and inhibiting airway differentiation came from an *NKX2.1* knock-down experiment in the Lin^POS^ organoids. A small decrease in *NKX2.1* expression was sufficient to decrease AT2-specific gene expression and increase *TP63* (Fig. 5L).

### Organoid assays can be used to predict the effects of human genetic variation

Overexpression of NKX2.1 containing a deleted DNA binding homeodomain showed that the homeodomain is essential for AT2 differentiation (Fig. 6A,B). Numerous naturally-occurring human variants in the NKX2.1 homeodomain have been described^12–14^. Many of these are associated with acute respiratory failure, others are predicted to be pathogenic. We hypothesized that the NKX2.1 OE assay would be a simple method to determine the effects of these variants on AT2-specific gene expression (Fig. 6C). The variants differentially affected organoid morphology, AT2 gene transcription and surfactant protein production (Fig. 6D-F), with the predicted pathogenic c.485_487 deletion behaving indistinguishably to the wildtype control. Interestingly, expression of the NKX2.1 variants (c.523G>T and c532C>T) often resulted in the production of mis-localised pro-SFTPC which was not processed to the mature form (Fig. 6D,F). This strongly suggests that NKX2.1 promotes multiple aspects of AT2 differentiation, not simply *SFTPC* transcription.

**Fig. 6.**
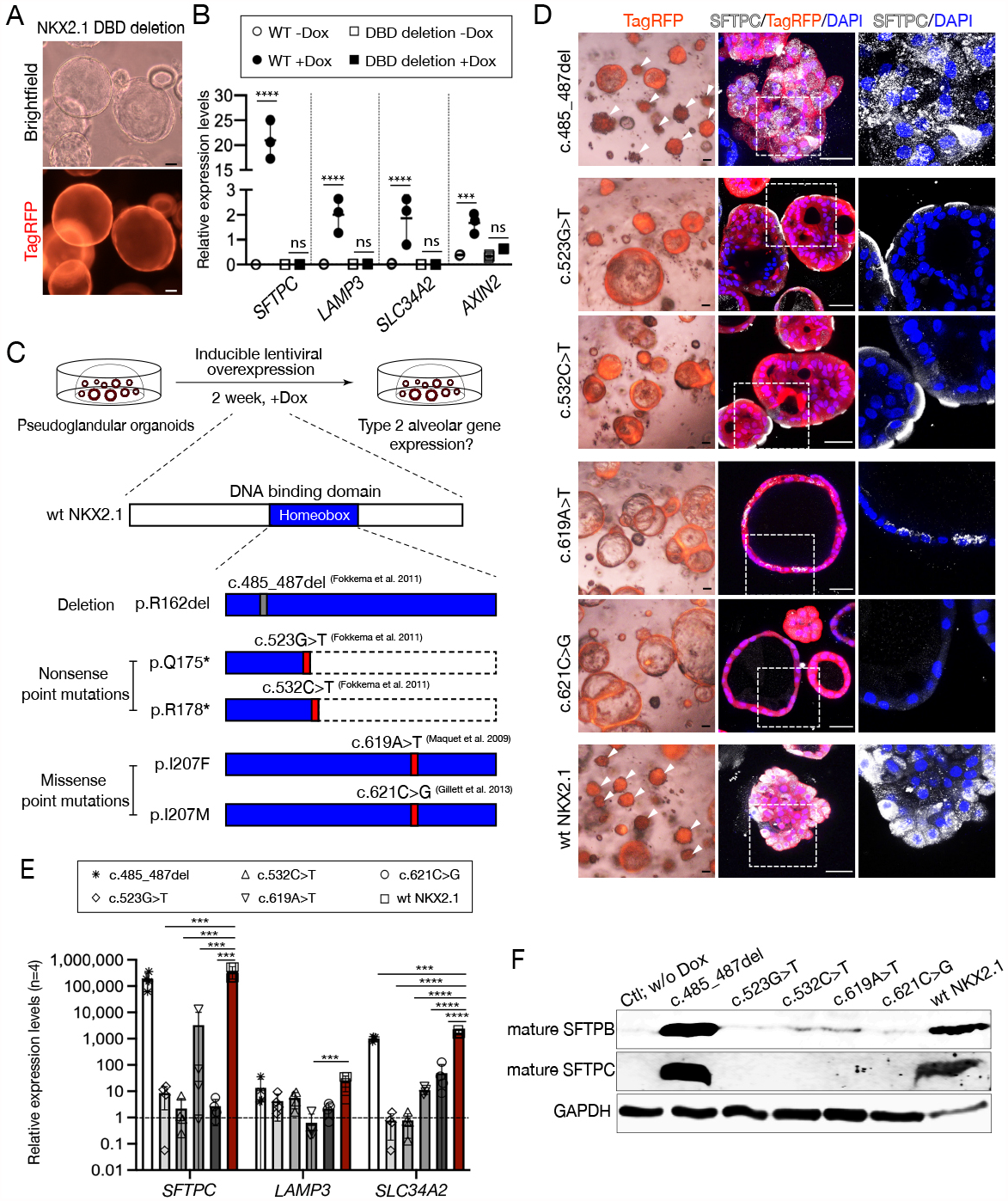
Analysis of naturally occurring human genetic variation using organoid assays. (A and B) Morphology (A) and gene expression profile (B) in the pseudoglandular organoids overexpressing wildtype NKX2.1, or a NKX2.1 lacking a DNA binding domain (DBD deletion), cultured for 2 weeks in the presence, or absence, of doxycycline (±DOX). (C) Diagram describing overexpression of wild type and mutant forms of NKX2.1 in the pseudoglandular organoids using doxycycline inducible lentiviral system. Individually, five different mutations were introduced into the DNA-binding homeobox domain; deletion of Arg^162^ (R162del)^14^, two nonsense point mutations (Q175*, R178*)^14^, and two missense point mutations (I207F, I207M)^12,13^ were tested. (D-F) Morphology and immunostaining (D), qRT-PCR (E), and western blot (F) analysis of the pseudoglandular organoids following overexpression of wildtype or mutant human NKX2.1 for 1 week. Data were normalized to doxycycline-non-treated lines; mean ± SD of 4 biological replicates. Significance was evaluated by 1-way ANOVA with Tukey multiple comparison post-test; \**P*<0.05, \*\**P*<0.01, \*\*\**P*<0.001. Western blot showing mature SFTPB and SFTPC. GAPDH was used for a loading control. DAPI indicates nuclei. Scale bar, 50 μm.

### Cultured canalicular stage tip cells differentiate readily into alveolar type 2 cells

NKX2-1 OE pseudoglandular organoids had higher levels of SFTPC and ACE2 than Lin^POS^ organoids, consistent with differentiation to AT2 fate (Fig. 7A; Extended Data Fig. 8A). We hypothesized that a medium change would allow passaged Lin^POS^ organoids to differentiate into AT2 cells. In medium containing DAPT (Notch inhibition), DCI (dexamethasone, cAMP, IBMX), CHIR (Wnt agonist) and SB431542 (TGFβ inhibition), *NKX2-1, SFTPC* and *ACE2* were upregulated and *SOX9, SOX2* and *TP63* downregulated (Fig. 7B,C). Moreover, the *SFTPC*-GFP reporter and the AT2-specific proteins LAMP3, HOPX and ACE2 were increased (Fig. 7C,D; Extended Data Fig. 8B-D). The pseudoglandular organoids did not differentiate towards AT2 fate in response to the same medium (Fig. 7E), confirming that the canalicular stage tips are in a distinct differentiation-ready state.

**Fig. 7.**
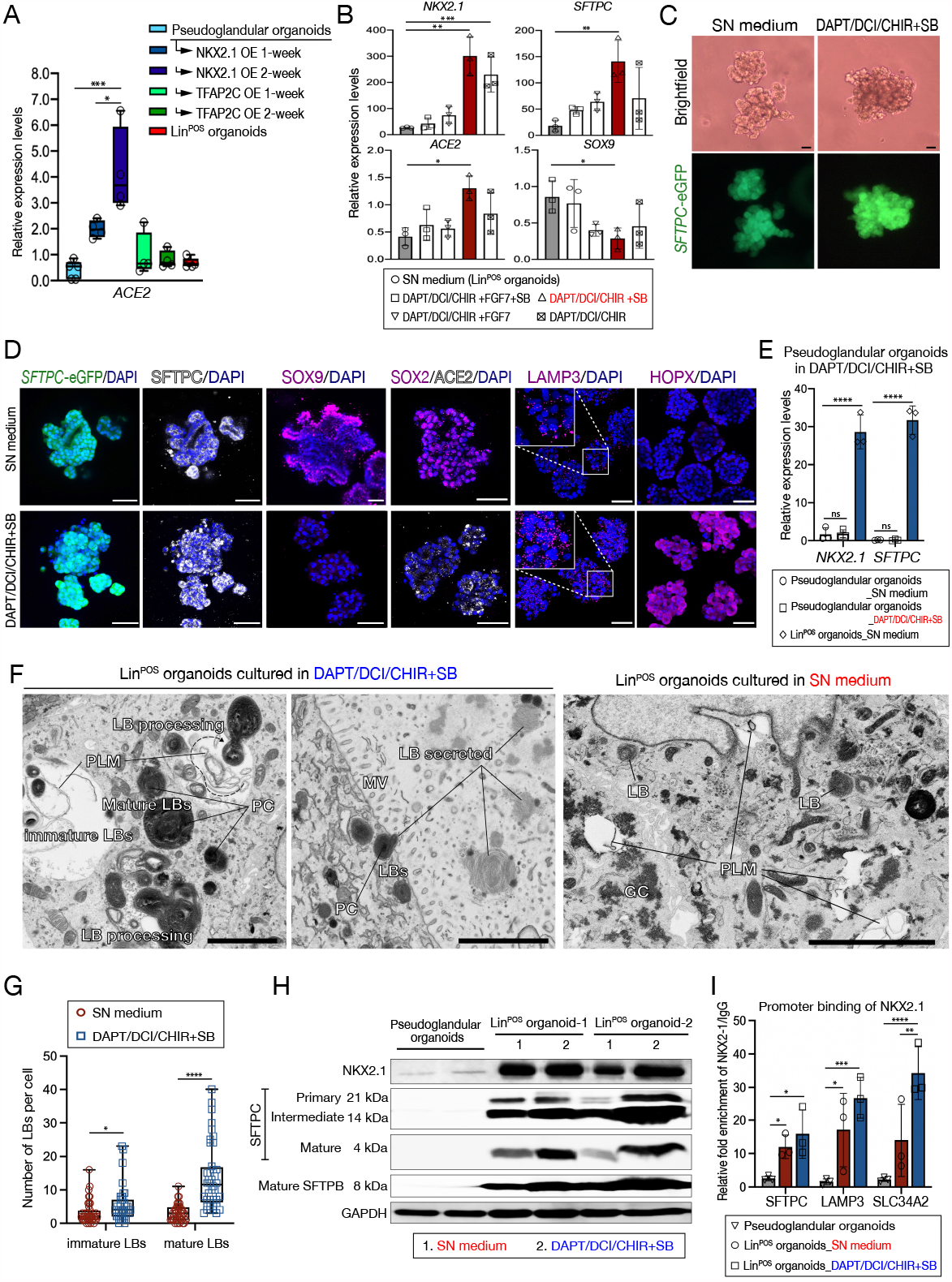
Efficient *in vitro* differentiation of Lin^POS^ organoids to alveolar type 2 cell fate. (A) qRT-PCR for *ACE2* in the pseudoglandular organoids overexpressing NKX2.1 or TFAP2C for 1 or 2 weeks. Data was normalized to fresh EPCAM^+^ cells from 20 pcw tip tissues; mean ± SD of four biological replicates. Significance was evaluated by 1-way ANOVA with Tukey multiple comparison post-test; \**P*<0.05, \*\**P*<0.01, \*\*\**P*<0.001. (B) qRT-PCR of the Lin^POS^ organoids cultured in SN medium or alveolar-induction culture conditions containing combinations of DAPT, <u>D</u>examethasone/<u>c</u>yclic AMP/ IBMX (DCI), CHIR with/without SB431542 (SB) and FGF7, for 1 week. *NKX2.1, SFTPC*, and *SOX9* levels were normalized to EPCAM^+^ cells, and *ACE2* was normalized to EPCAM^-^ cells, freshly sorted from 20 pcw tip tissues; mean ± SD of four biological replicates. Significance was evaluated by 1-way ANOVA with Dunnett multiple comparison post-test; \**P*<0.05, \*\**P*<0.01, \*\*\**P*<0.001. (C) Morphology and fluorescent images of the Lin^POS^ organoids cultured in DAPT/DCI/CHIR with SB, or in control SN medium, for 1 week. (D) Immunofluorescent analysis of *SFTPC*-GFP^+^ Lin^POS^ organoids cultured in DAPT/DCI/CHIR with SB or in the SN medium for 1 week. DAPI indicates nuclei. Scale bar, 50 μm. (E) qRT-PCR of *NKX2.1* and *SFTPC* in the pseudoglandular organoids cultured in the SN medium or in DAPT/DCI/CHIR plus SB. Data were normalized to EPCAM^+^ cells freshly isolated from 20 pcw tip tissues; mean ± SD of three biological replicates. Significance was evaluated by 1-way ANOVA with Tukey multiple comparison post-test; \*\*\*\**P*<0.0001. (F) Electron microscopy images of Lin^POS^ organoids cultured in DAPT/DCI/CHIR with SB (*left*) or SN medium (*right*). LBs, lamellar bodies; PC, projection core; MV, microvilli; GC, glycogen; PLM, Primitive lipid membrane within a pool of monoparticulate glycogen at an early stage in the formation of LBs. Scale bar, 3 μm. (G) Numbers of LBs per cells in Lin^POS^ organoids cultured in SN medium (*red*) or DAPT/DCI/CHIR with SB (*blue*). Immature and mature LBs were measured in total 40 cells from two biological samples for each condition. Significance evaluated by unpaired student *t*-test; \**P*<0.05, \*\**P*<0.01, \*\*\**P*<0.001, \*\*\*\**P*<0.0001. (H) Western blot showing NKX2.1 levels and SFTPB/SFTPC processing in the Lin^POS^ organoids cultured in the SN medium, or in DAPT/DCI/CHIR/SB and the pseudoglandular organoids cultured in the SN medium for 1 week. GAPDH was used for a loading control. (I) Chromatin immunoprecipitation (ChIP)-qPCR analysis for quantifying relative enrichment of NKX2.1 binding on the promoter regions of type 2 alveolar lineage markers, *SFTPC, LAMP3*, and *SLC34A2* in the organoids cultured in the SN medium, or in DAPT/DCI/CHIR plus SB. Data was normalized to the IgG control; mean ± SD of three biological replicates.

Electron microscopy revealed that the Lin^POS^ organoids in the SN medium contained rare, immature lamellar bodies usually surrounded by glycogen (Fig. 7F,G). Whereas higher numbers of lamellar bodies with a characteristic surfactant projection core were readily visible in the differentiated organoids (Fig. 7F,G)^15,16^. NKX2.1 protein levels were increased following AT2 differentiation (Fig. 7H). The differentiated cells also more efficiently processed pro-SFTPC and SFTPB to the mature form (Fig. 7H)^16^. Moreover, NKX2.1 binds more strongly to the promoters of AT2-specific genes following differentiation (Fig. 7I). These data demonstrate that the Lin^POS^ organoids are readily differentiated to an AT2-like fate and confirm the importance of NKX2.1 levels in this process.

## DISCUSSION

We show that the distal tip cells at the canalicular stage of human lung development retain their progenitor status yet exhibit aspects of AT2 gene expression and can be isolated using specific surface proteins. Late-tip cell (Lin^POS^) organoids self-renew, capture features of the canalicular stage of human lung development, have extensive open chromatin and can be readily differentiated to AT2-like cells. We have used this organoid system to demonstrate that Wnt signalling and NKX2.1 are required for human AT2 cell differentiation. Additionally, we show that antagonistic signalling interactions between differentiating fibroblasts and myofibroblasts provide a spatial component to Wnt activation allowing patterning of the alveolar epithelium into specific lineages. We also demonstrate that our organoid system can be used to study human genetic variation.

One of major insights of this work is the demonstration that the molecular acquisition of alveolar features precedes morphological changes occurring in the tip epithelium at the canalicular stage. Our Lin^POS^ organoids are derived from the CD44^+^,CD36^+^ canalicular stage tips. The CD44^+^,CD36^+^ cells are located at the tips of the organoids, maintain expression of SOX9 and AT2 markers, self-renew and give rise to airway-fated cells in the centre of the organoids (Fig. 2). When provided with appropriate cues they differentiate to an AT2-like cell (Fig. 7). *In vivo*, tip cells acquire AT2 markers gradually between 13 and 15 pcw (Fig. 1). We hypothesize that during this transition period (∼13–15 pcw) the tips are generating the final branch of the airway epithelium and at ∼15 pcw switch to generating alveolar fated daughter cells. However, due to well-documented tip progenitor cell plasticity in transplantation assays^17,18^, it is not yet possible to test this definitively.

We identify Wnt signalling as a key driver of human AT2 fate and patterning. Wnt signalling promotes AT2 differentiation of the tip epithelium *in vitro* and *in vivo* (Fig. 3,4). This is consistent with previous reports in mouse^19^. Similarly, human NKX2.1^+^ lung progenitors derived from PSCs expressed alveolar epithelial markers in response to Wnt^5^. Our experiments with primary tissue support a model in which opposing signals from differentiating alveolar fibroblasts and myofibroblasts spatially restrict late tip and AT2 identity in the canalicular stage human lung (Fig. 4). These data are analogous to a recent mouse report where developing AT1 cells are aligned with, and signal to, differentiating myofibroblasts^20^. It will be interesting to test in the future whether myofibroblast inhibition of AT2 cell fate occurs in pulmonary fibrosis where myofibroblasts are expanded and AT2 cells lost.

We clearly demonstrate that NKX2.1 is a key upstream TF driving the onset of the alveolar program whilst supressing the airway program (Fig. 5), consistent with reported roles in lung cancers^21^. Our data also strongly suggest that Wnt is upstream of NKX2.1 during alveolar differentiation. By contrast, FGF signalling and TFAP2C cooperate to promote airway fate. Ectopic expression of TFAP2C promoted the expression of basal cell markers, but not other airway lineages (Fig. 5). This could mean the culture conditions are permissive only for basal cell differentiation. Alternatively, TFAP2C may be specific for basal cell specification. The latter interpretation would be consistent with a report that TFAP2C activates TP63 expression during epidermal lineage maturation^22^.

In summary, we have identified a distinct late-tip progenitor cell state in the developing human lung. Culture of these cells as organoids has allowed us to investigate the roles of Wnt signalling and NXKX2.1 in human AT2 cell development. Moreover, this differentiating organoid system will be useful for understanding the next phases of the alveolar maturation process at the saccular and alveolar stages in the developing human lung.

## ACKNOWLEDGEMENTS

We would like to acknowledge the Gurdon Institute Imaging Facility and Dr Karin Mueller of Cambridge Advanced Imaging Centre for microscopy support. Prof Azim Surani for epigenetics advice. KL is supported by Basic Science Research Program through the National Research Foundation of Korea (NRF) funded by the Ministry of Education (2018R1A6A3A03012122). DS is supported by a Wellcome Trust PhD studentship (109146/Z/15/Z) and the Department of Pathology, University of Cambridge ELR. is supported by Medical Research Council (MR/P009581/1; MR/S035907/1). Gurdon Institute Core support from the Wellcome Trust (203144/Z/16/Z) and Cancer Research UK (C6946/A24843). KBM and SAT acknowledge funding from the MRC (MR/S035907/1) and from Wellcome (WT211276/Z/18/Z and Sanger core grant WT206194).

## AUTHOR CONTRIBUTIONS

Conceptualization, KL and ELR; Methodology, Investigation, and Validation, KL, DS ; Software and Formal Analysis, KL, WT, PH; Writing–Original Draft by KL; Writing – Review & Editing, ELR; Funding Acquisition and Supervision: ELR, SAT, JCM, KBM.

## FIGURE LEGENDS

**Extended Data Fig. 1.**
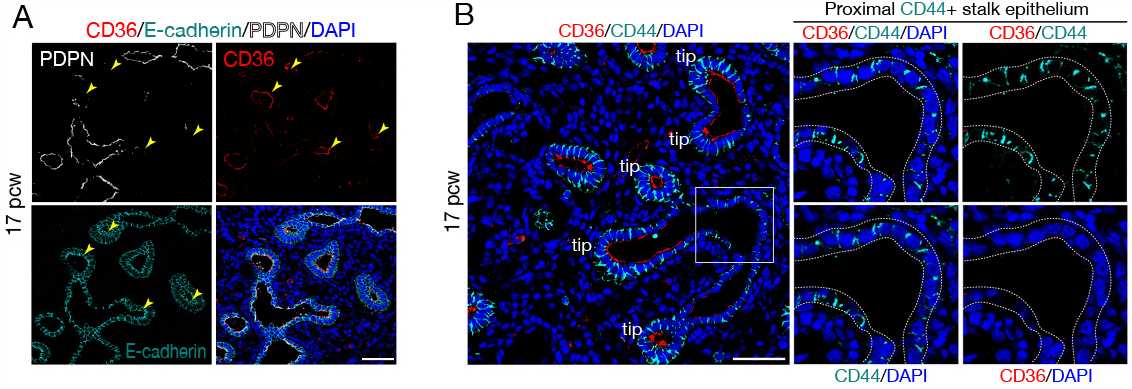
Characterization of the lung tip epithelium at the canalicular stages. (A and B) Frozen sections of human fetal lung tissues at 17 pcw. Stained for CD36, E-cadherin and PDPN (A) and CD36, CD44 (B). Arrowheads indicate CD36^+^PDPN^-^ tips. Inset (B) shows a CD44+, CD36- stalk epithelial region. DAPI, nuclei. Scale bars, 50 μm.

**Extended Data Fig. 2.**
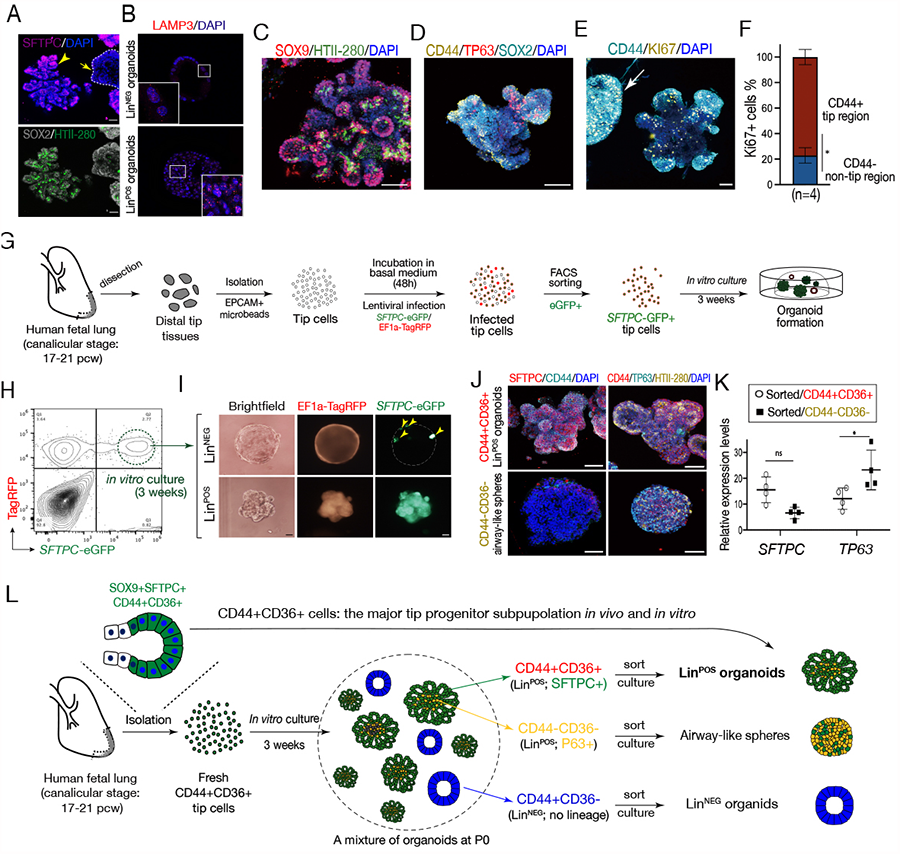
Characterization of the canalicular stage lung tip organoids. (A and B) Immunofluorescence analysis of the Lin^POS^ (arrowheads) and Lin^NEG^ organoids (arrow) at passage 1 cultured in the self-renewing medium, showing the alveolar lineage markers, SFTPC, HTII-280 (A) and LAMP3 (B, *lower* panel) were expressed in the Lin^POS^ organoids, but not in the Lin^NEG^ organoids. DAPI, nuclei. Scale bar, 50 μm. (C-E) Immunofluorescence images (C-E) of the Lin^NEG^ and Lin^POS^ organoids originating from sorted CD44^+^CD36^+^ tip epithelium from 20 pcw lung. Antibodies against SOX9, HTII-280 (C), CD44, TP63, SOX2 (D) and CD44, KI67 (E). Arrow (E) indicates a Lin^NEG^ organoid. DAPI indicates nuclei. Scale bar, 50 μm. (F) The percentage of KI67^+^ cells in the CD44^+^ tip and CD44^-^ non-tip regions are represented as mean ± SD of biological 4 replicates. (G-I) Diagram (G) illustrating isolation and *in vitro* culture of the tip epithelial cells infected with lentivirus harbouring *SFTPC* promoter-driven eGFP and EF1a promoter-driven TagRFP. The cells expressing *SFTPC-eGFP* were sorted at 48h post-infection and cultured in the self-renewing medium for 3 weeks (H). The Lin^NEG^ and Lin^POS^ organoids at passage zero derived from the *SFTPC-eGFP* positive-sorted tip epithelial cells (I). Low eGFP signals remained in some Lin^NEG^ organoids (arrowhead) confirming their derivation from SFTPC^+^ tip cells. Scale bar, 50 μm. (J and K) Expression of lineage markers was investigated by immunostaining (J) and qRT-PCR (K) in the Lin^POS^ organoids and airway-like spheres at passage 1 derived from the CD44^+^CD36^+^ or CD44^-^CD36^-^ passage zero subpopulations respectively. Data was normalized to EPCAM^+^ cells freshly sorted from 20 pcw tip tissues and represented as mean ± SD of 4 biological replicates. Significance was evaluated by 2-way ANOVA with Bonferroni multiple comparison post-test; ns: not significant, \**P*<0.05. (L) Diagram summarising the organoid experiments performed in Fig. 2. CD44^+^CD36^+^ cells from canalicular stage lung tissues are the major tip progenitor subpopulation *in vitro*, growing into self-renewing Lin^POS^ organoids showing key features of the canalicular stage lung tip cells.

**Extended Data Fig. 3.**
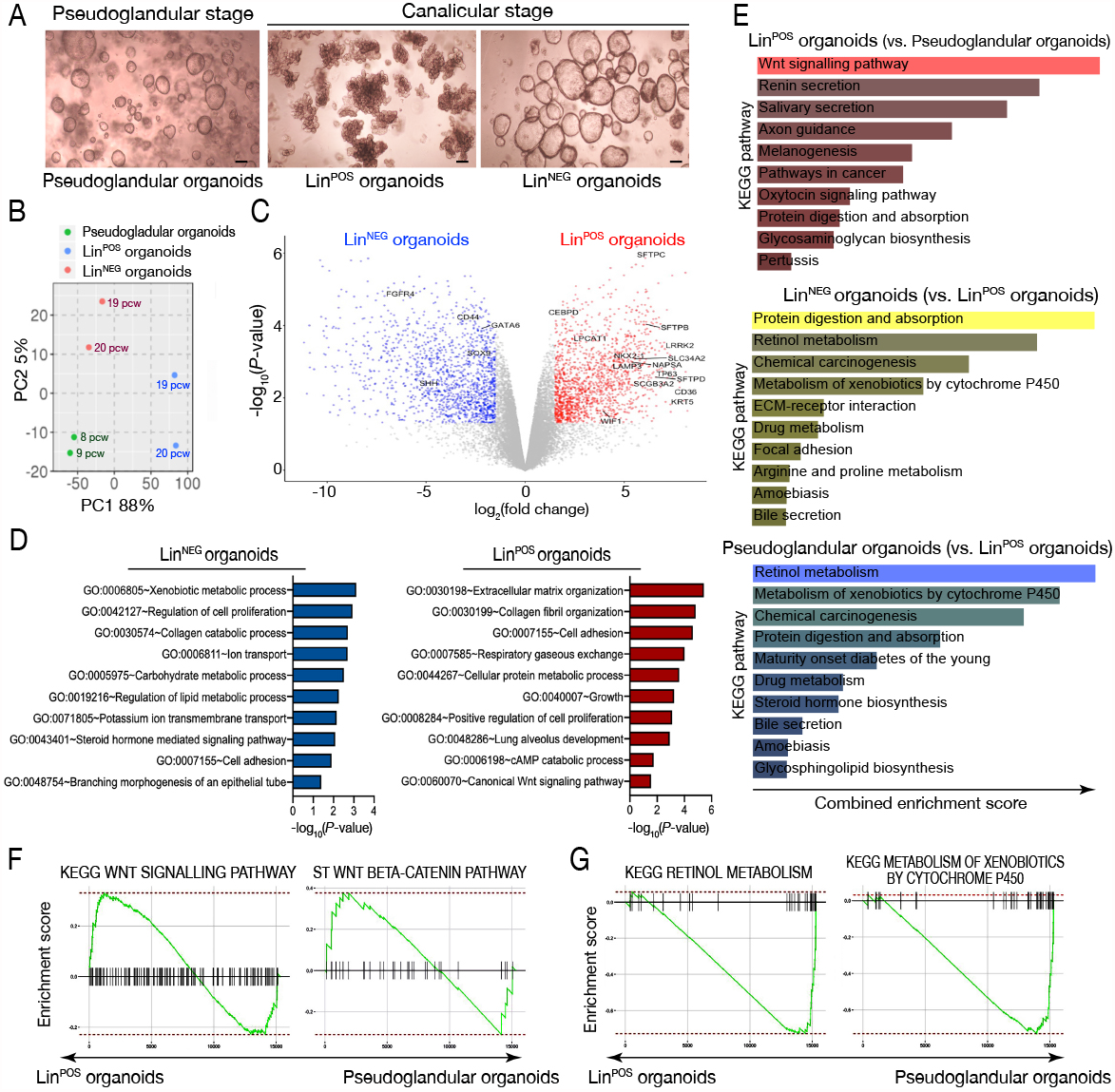
Transcriptomic analysis of the lung tip organoids. (A) Morphology of the pseudoglandular organoids from the pseudoglandular stage, and the Lin^POS^ and Lin^NEG^ organoids from the canalicular stage. The Lin^POS^ and Lin^NEG^ organoids grown from EPCAM^+^ tip epithelial cells were manually separated and cultured in the self-renewing culture condition. (B) Principal component analysis of bulk-RNA seq data using pseudoglandular organoids, Lin^POS^ and Lin^NEG^ organoids. (C) Volcano plot showing differentially expressed genes between Lin^POS^ organoids (*red*) versus Lin^NEG^ organoids (*dark blue*); log_2_FC > 4. (D) Gene ontology (GO) enrichment analysis performed for biological process (BP)-associated GO terms on the differentially expressed genes between the Lin^POS^ and Lin^NEG^ organoids; log_2_FC > 4. (E) KEGG pathway analysis using Enrichr. Length of coloured bars indicates combined enrichment score by adjusted p-value < 0.05. (F and G) Gene set enrichment (GSEA) analysis of the differentially expressed genes of the Lin^POS^ organoids (F) and pseudoglandular organoids (G).

**Extended Data Fig. 4.**
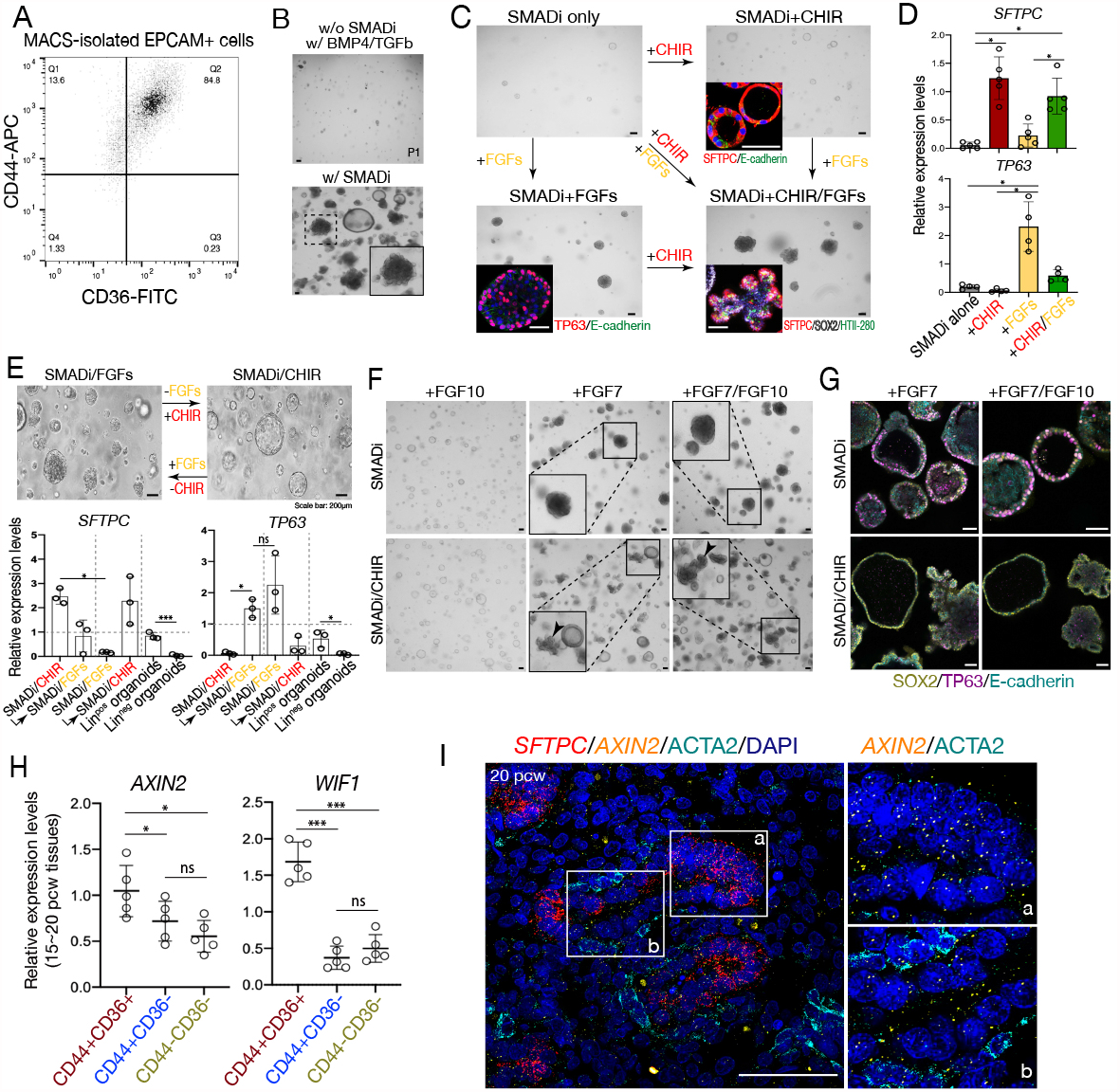
High plasticity of the canalicular stage tip epithelial cells in response to Wnt and FGF signalling. (A) The population of EPCAM^+^ tip epithelial cells in the distal lung tissue at 20 pcw was isolated using MACS and were analysed by CD44 and CD36 expression using FACS. (B) Tip epithelial cells cultured with, or without, SMAD inhibition for 3 weeks. The self-renew medium condition (w/ SMADi; *lower* panel) was used for positive control. Scale bar, 200 μm. (C, D) After 2 weeks growing in SMADi conditions, cells were transferred to culture medium containing CHIR, or FGFs, or CHIR/FGFs. After 2 weeks of exposure to the different culture conditions, the epithelial organoids were stained with lineage makers including SFTPC or TP63 (C) and relative mRNA levels of *SFTPC* and TP63 were measured by qRT-PCR (D). Data were normalized to the Lin^POS^ organoids; mean ± SD of at least 4 biological replicates. Significance was evaluated by 1-way ANOVA with Tukey multiple comparison post-test; ns: not significant, \**P*<0.05, \*\**P*<0.01 and \*\*\**P*<0.001. Scale bar, 200 μm. (E) After 2 weeks growing in SMADi/FGFs (or SMADi/CHIR) the organoids were transferred to SMADi/CHIR (or SMADi/FGFs) for another 2 weeks. After 2 weeks of exposure to the different culture conditions organoid morphology altered and qRT-PCR for lineage markers was performed. Data were normalized to the Lin^POS^ organoids; mean ± SD of 3 biological replicates. Significance was evaluated by 1-way ANOVA with Tukey multiple comparison post-test; ns: not significant, \**P*<0.05, \*\**P*<0.01 and \*\*\**P*<0.001. Scale bar, 200 μm. (F and G) Tip epithelial cells cultured with FGF7, FGF10, or FGF7 and FGF10 (F); scale bar, 200 μm. After 2 weeks of exposure to each culture condition, the organoids were immunostained for SOX2, TP63 and E-cadherin (G); scale bar, 50 μm. (H) Gene expression profile of the freshly isolated lung epithelial cells from the canalicular stage human lung tissues sorted by CD44^+^CD36^+^, CD44^+^CD36^-^ and CD44^-^CD36^-^. Data normalized to freshly isolated EPCAM^+^ cells from 20 pcw tip tissues; mean ± SD of 5 biological replicates aged from 15∼20 pcw. Significance was evaluated by 1-way ANOVA with Tukey multiple comparison post-test; ns: not significant, \**P*<0.05, \*\**P*<0.01 and \*\*\**P*<0.001. (I) Frozen sections of human fetal lung tissues at 20 pcw were stained for ACTA2 followed by *in situ* HCR for *SFTPC* and *AXIN2*.

**Extended Data Fig. 5.**
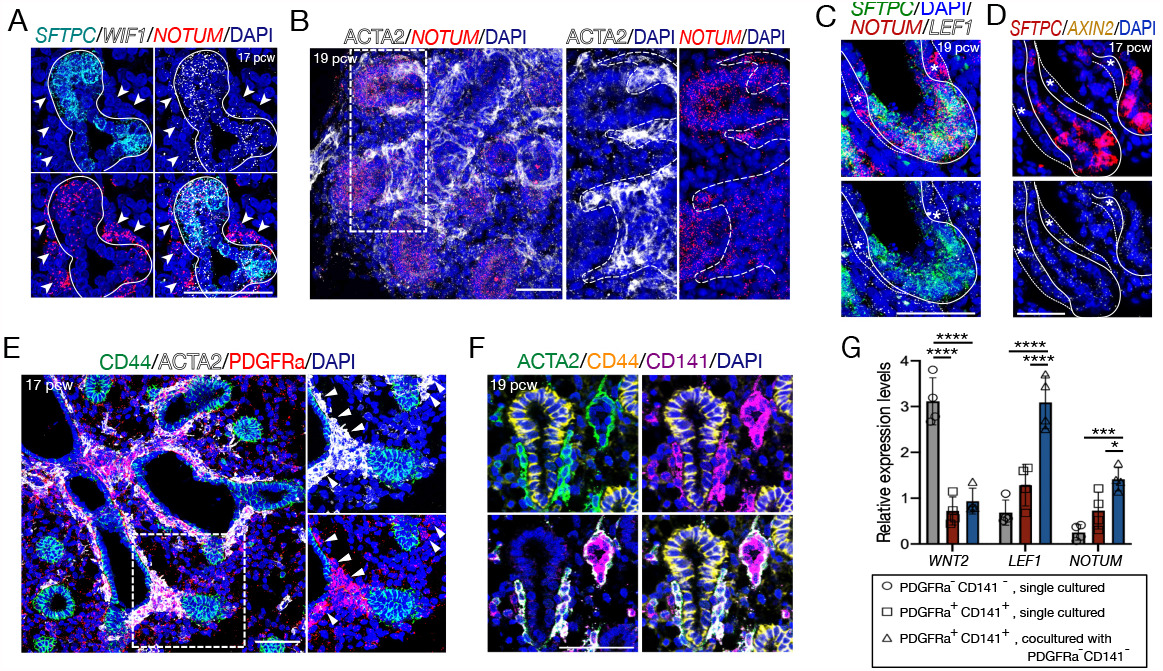
Wnt-responsive NOTUM^+^ myofibroblasts in the distal human fetal lung. (A-F) Human fetal lung sections at 17 (A, D, E) and 19 pcw (B, C, F). A. *SFTPC, WIF1, NOTUM*. B. ACTA2, *NOTUM*. C. *NOTUM, LEF1*. D. *SFTPC, AXIN2*. E. CD44, ACTA2, PDGFRA. F. ACTA2, CD44, CD141. Arrows and asterisks indicate ACTA2^+^ PDGFRA^+^ *NOTUM*^+^ myofibroblasts. Lines and dashed lines indicate the boundaries of epithelial cells and myofibroblasts, respectively. (G) Relative mRNA levels from myofibroblasts and alveolar fibroblasts cultured alone, or cocultured in transwells. Data were normalized to the whole freshly isolated lung fibroblast population; mean ± SD of biological 4 replicates. Significance was evaluated by unpaired student *t*-test; \**P*<0.05, \*\**P*<0.01, \*\*\**P*<0.001. DAPI indicates nuclei. Scale bar, 50 μm.

**Extended Data Fig. 6.**
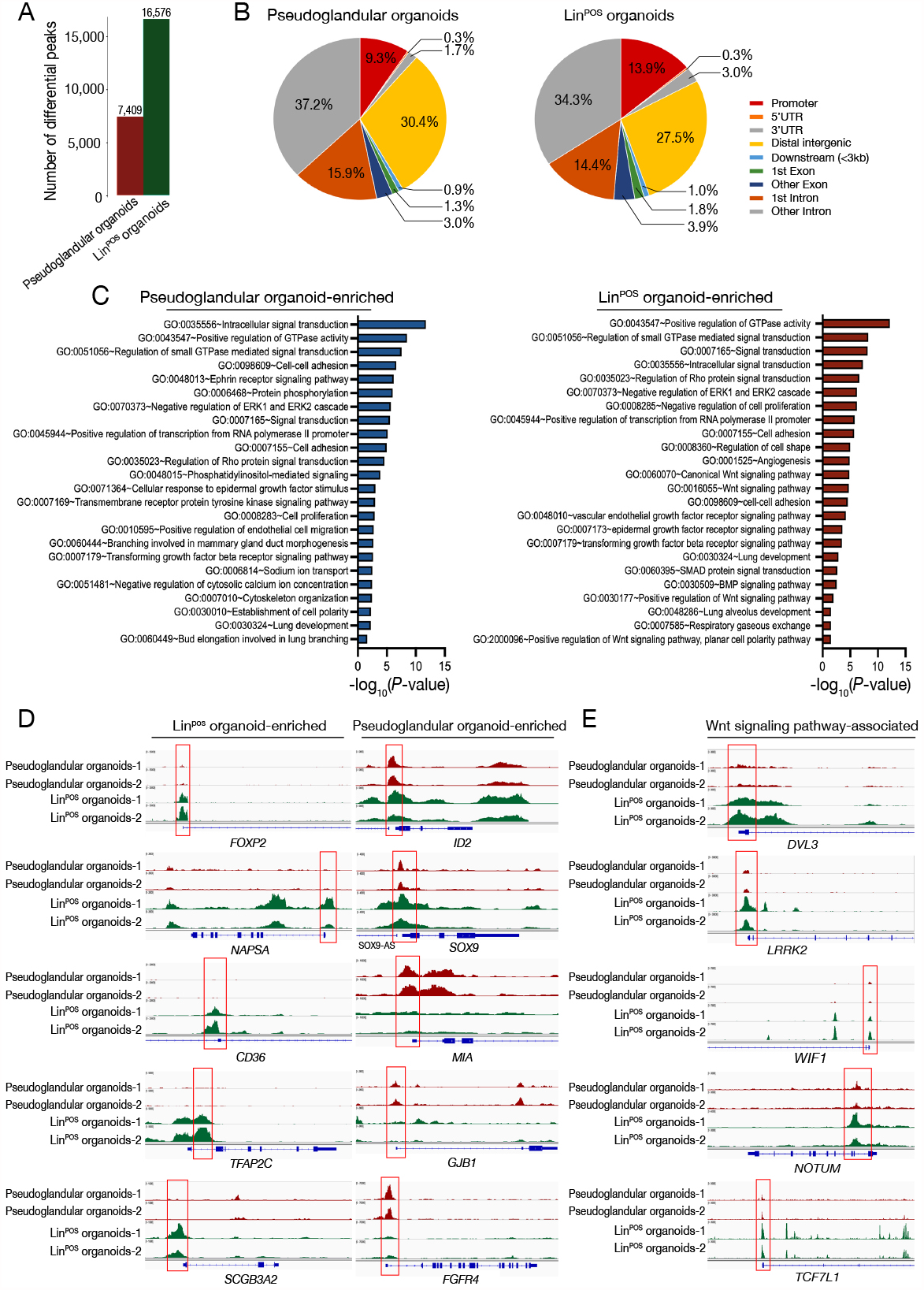
Chromatin accessibility analysis of the pseudoglandular and Lin^POS^ organoids. (A) Analysis of chromatin accessibility in the pseudoglandular and Lin^POS^ organoids by bulk-ATAC seq. The number of differential accessible chromatin regions which differ between the organoids was identified; fold change > 2 and FDR< 0.05. (B) Pie charts representing the genomic distribution of global accessible chromatin regions in pseudoglandular organoids and Lin^POS^ organoids. (C) Biological Process-associated GO term analysis using the differential accessible chromatin regions highly enriched in the pseudoglandular organoids and Lin^POS^ organoids. (D and E) IGV image shots of representative ATAC seq tracks at loci showing differential accessible chromatin regions between pseudoglandular organoids and Lin^POS^ organoids. Red box indicates the promoter regions.

**Extended Data Fig. 7.**
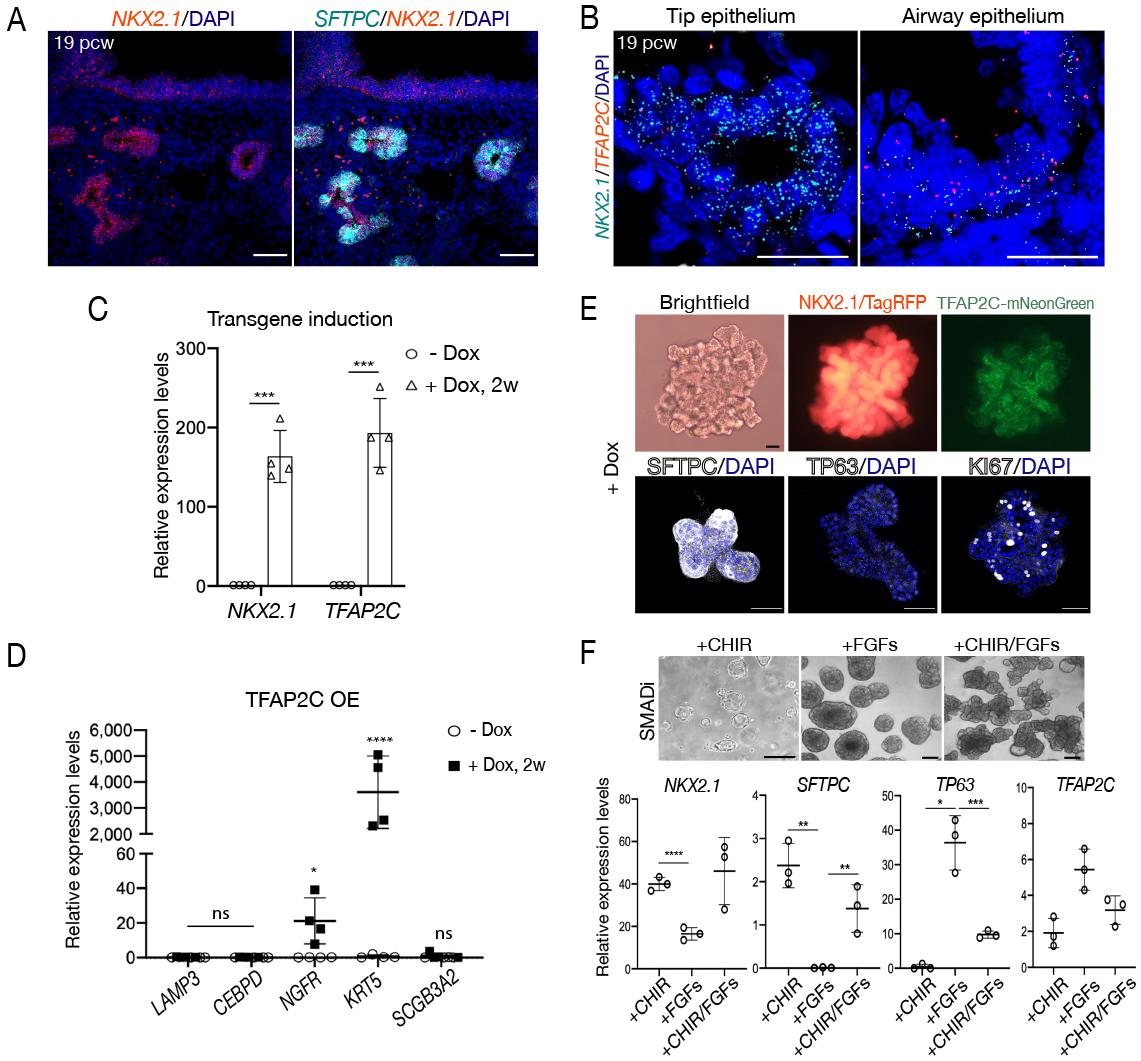
NKX2.1 drives the onset of the alveolar program whilst supressing the airway program. (A and B) *In situ* HCR images for detecting transcripts, *SFTPC* or *TFAP2C* with *NKX2.1* at 19 pcw. (C) Transgene induction following doxycycline treatment for 2 weeks measured by qRT-PCR. Data was normalized to the untreated group; mean ± SD of four biological replicates. Significance was evaluated by 1-way ANOVA with Tukey multiple comparison post-test; ns: not significant, \*\*\*\**P*<0.0001. (D) qRT-PCR of the pseudoglandular organoids overexpressing TFAP2C for 2 weeks. Data was normalized to EPCAM^+^ positive cells freshly isolated from 20 pcw tip tissues; mean ± SD of biological 4 replicates. Significance was evaluated by 1-way ANOVA with Tukey multiple comparison post-test; ns: not significant, \**P*<0.05, \*\**P*<0.01, \*\*\**P*<0.001 and \*\*\*\**P*<0.0001. (E) Morphology and fluorescent images of pseudoglandular organoids overexpressing both NKX2.1 and TFAP2C for 2 weeks. (F) qRT-PCR of endogenous *NKX2.1, SFTPC*, and *TP63* in the Lin^POS^ organoids cultured in medium containing CHIR, FGF7 or CHIR/FGF7. Data was normalized to EPCAM^+^ cells freshly isolated from 20 pcw tip tissues; mean ± SD of three biological replicates. Significance was evaluated by 1-way ANOVA with Tukey multiple comparison post-test; ns: not significant, \**P*<0.05, \*\**P*<0.01, \*\*\**P*<0.001, \*\*\*\**P*<0.0001. DAPI indicates nuclei. Scale bar, 50 μm.

**Extended Data Fig. 8.**
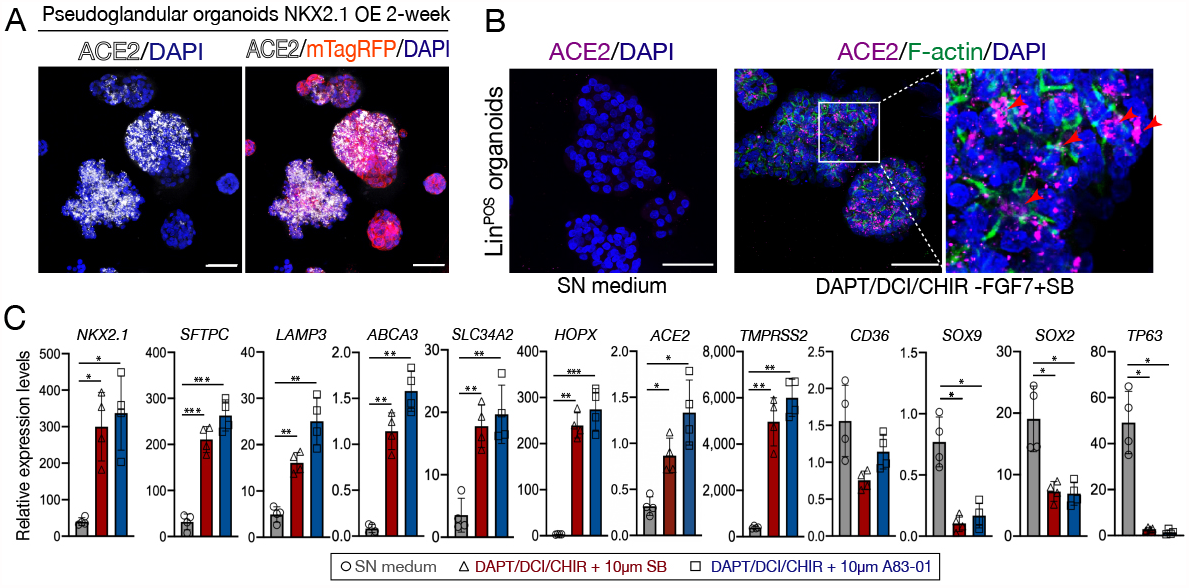
Type 2 alveolar differentiation of the cultured canalicular stage tip cells. (A and B) Immunofluorescent analysis of ACE2 in (A) pseudoglandular organoids overexpressing NKX2.1 and (B) Lin^POS^ organoids were cultured in DAPT/DCI/CHIR/SB or in the SN medium for 1 week. Phalloidin (F-actin) marks apical membrane of epithelial cells in the organoids. Scale bar, 50 μm. (C) qRT-PCR of Lin^POS^ organoids cultured in the SN medium or in DAPT/DCI/CHIR with SB, or A83-01, for 1 week. Data were normalized to EPCAM^+^ cells freshly isolated from 20 pcw tip tissues; mean ± SD of four biological replicates. Significance was evaluated by 1-way ANOVA with Tukey multiple comparison post-test; \**P*<0.05, \*\**P*<0.01, \*\*\**P*<0.001.

## MATERIALS and METHODS

### Human embryonic and foetal lung tissue

Human embryonic and foetal lung tissues were provided from terminations of pregnancy from Cambridge University Hospitals NHS Foundation Trust under permission from NHS Research Ethical Committee (96/085) and the MRC/Wellcome Trust Human Developmental Biology Resource (London and Newcastle, University College London (UCL) site REC reference: 18/LO/0822; Newcastle site REC reference: 18/NE/0290; Project 200454; www.hdbr.org). Sample age ranged from 4 to 23 weeks of gestation (post-conception weeks; pcw). Stages of the samples were determined according to their external physical appearance and measurements. All the samples used for the current study had no known genetic abnormalities.

### *In vitro* culture of human fetal lung organoids

The isolated tip epithelial cells were embedded in Matrigel (Corning, 356231) and cultured in 48-well plates in self-renewal (SN) medium: Advanced DMEM/F12 supplemented with 1x GlutaMax, 1 mM HEPES and Penicillin/Streptomycin, 1X B27 supplement (without Vitamin A), 1X N2 supplement, 1.25 mM n-Acetylcysteine, 50 ng/ml recombinant human EGF (PeproTech, AF-100-15), 100 ng/ml recombinant human Noggin (PeproTech, 120-10C), 100 ng/ml recombinant human FGF10 (PeproTech, 100-26), 100 ng/ml recombinant human FGF7 (PeproTech, 100-19), 3 μM CHIR99021 (Stem Cell Institute, University of Cambridge) and 10 mM SB431542 (Bio-Techne, 1614). The culture medium was replaced every 2 days and the organoids were usually split 1:3 once per week by breaking them into small fragments. Numbers of replicates are indicated in figure legends. 10 ng/μl recombinant human BMP4 (Peprotech, 120-05) and 10 ng/μl recombinant human TGF-β1 (Peprotech, 100-21) were added to the medium instead of adding SB431542 and Noggin to activate dual SMAD signalling (Extended Data Fig. 4B).

To perform *in vitro* co-culture experiments, freshly sorted 2×10^5^ PDGFRA^+^CD141^+^ myofibroblasts or PDGFRA^-^CD141^-^ fibroblasts were mixed with *SFTPC*-eGFP^+^ Lin^POS^ organoids in 100 μl Matrigel and then loaded into an insert of transwell (Merck, CLS3493). On the bottom well plates, coated with Collagen (Merck, CLS3493), 4×10^5^ PDGFRA^-^CD141^-^ fibroblasts were plated in the culture medium containing 2% fetal bovine serum (FBS; Thermo, 10500064) in the Advanced DMEM/F12 supplemented with 1x GlutaMax, 1 mM HEPES and Penicillin/Streptomycin. 100 μl culture medium was added every 2 days. For coculture of PDGFRA^+^CD141^+^ myofibroblasts with PDGFRA^-^CD141^-^ fibroblasts, 2×10^5^ of PDGFRA^+^CD141^+^ myofibroblasts were plated on the insert and 4×10^5^ of PDGFRA^-^CD141^-^ fibroblasts were plated on the bottom well plate. After 2 weeks of co-cultures the organoids, or the mesenchymal cells, were harvested for further analysis.

### Isolation of tip epithelial cells, myofibroblasts, and alveolar fibroblasts

For isolation of CD44^+^, or CD44^+^CD36^+^, tip epithelium directly from the distal lung tissues, the tissues were finely dissected into tiny pieces and enzymatically digested into single cells by incubating them in a dissociation solution containing 0.125 mg/ml Collagenase (Merck, C9891), 1 U/ml Dispase (Thermo Fisher Scientific, 17105041) and 0.1 U/μl DNAase (Merck, D4527), in a rotating incubator for 1 h at 37°C. After rinsing in washing buffer containing 2% FBS in cold PBS the cells were filtered by 100 μm strainer and harvested by centrifugation. The cell pellets were resuspended and treated with RBC lysis buffer (BioLegend, 420301). Next, the cells were rinsed in the washing buffer and then incubated with primary antibodies against CD45 (1:100; PE-Cy7 conjugated, Thermo Fisher Scientific, 25-9459-42), CD31 (1:100; PE-Cy7 conjugated, Thermo Fisher Scientific, 25-0319-42), EPCAM (1:100; PE-conjugated; BioLegend, 324206), CD44 (1:200; APC-conjugated; BioLegend, 103012), and CD36 (1:100; FITC, conjugated; Thermo Fisher Scientific, 11-0369-42), with a viability dye, Zombie (Biolegend, 423113) for 25 min on ice. Following removal of dead cells and immune/endothelial cells, the EPCAM+ epithelial cells were sorted by CD44 and/or CD36 expression by FACS (BD Influx™ Cell Sorter) (Fig. 1G).

Alternatively, the tip epithelial cells were isolated by EPCAM^+^ magnetic-activated cell sorting (MACS) beads according to the manufacturer’s instruction (CD326 MicroBeads, human, Miltenyi Biotec) from the distal lung tissues. Then, the enriched EPCAM^+^ epithelial cells were sorted by CD44 and/or CD36 expression by FACS (SH800S Cell Sorter) to more purely enrich the tip cell population (Fig. 2A).

To purify myofibroblasts and alveolar fibroblasts, the single cells dissociated from the distal lung tissues were incubated with the following primary antibodies: CD45 (1:100; PE-Cy7 conjugated, Thermo Fisher Scientific, 25-9459-42), CD31 (1:100; PE-Cy7 conjugated, Thermo Fisher Scientific, 25-0319-42), CD9 (1:100; PE-Cy7 conjugated, BioLegend, 312115), EPCAM (1:100; FITC-conjugated, 324204), PDGFRA (1:100; APC-conjugated, BioLegend, 313511), CD141 (1:100; PE-conjugated, BioLegend, 344104), with the viability dye, Zombie (Biolegend, 423113). After removing dead cells, immune/endothelial cells, airway smooth muscle cells, and epithelial cells, the cells are sorted by PDGFRA and/or CD141 expression using BD Influx Cell Sorter. The sorted cells were directly applied to an organoid coculture or a gene expression analysis.

### Type 2 alveolar differentiation of Lin^POS^ organoids

The Lin^POS^ organoids were embedded in Matrigel and cultured for 7 days in alveolar type 2 (AT2) differentiation medium: Advanced DMEM/F12 supplemented with 1x GlutaMax, 1 mM HEPES and Penicillin/Streptomycin, 1X B27 supplement (without Vitamin A), 1x N2 supplement, 1.25 mM n-Acetylcysteine, 10mM CHIR99021, 50 μM Dexamethasone (Merck, D4902), 0.1 M 8-Bromoadenosine 3’5’-cyclic monophosphate (cAMP; Merck, B5386), 0.1 M 3-Isobutyl-1-methylxanthine (IBMX; Merck, 15679), 50 mM DAPT (Merck, D5942) with 10 mM SB431542 or 10 mM A83-01 (Tocris, 2939). The culture medium was replaced every 2 days without passaging.

### Lentiviral transduction

To introduce a reporter system into the tip epithelial cells, the lentiviral vector pHAGE hSPC-eGFP-W given from Darrell Kotton (Addgene plasmid # 36450; http://n2t.net/addgene:36450; RRID: Addgene_36450) was modified by inserting EF1a-promoter TagRFP cassette. The tip epithelial cells were infected with the modified lentiviral vector for 24 h at 37°C in a single cell suspension in the SN medium containing 10 μM Y-27632 (Merck, 688000). After 24 h, the cells were embedded to the Matrigel and cultured in the SN medium containing 10 μM Y-27632 for another 48 h to support single cell survival. The cultured cells were further sorted by eGFP/TagRFP signals to enrich the infected cells.

For overexpressing NKX2.1 and/or TFAP2C, Tet-ON 3G doxycycline (Dox)-inducible lentiviral vector (Takara, 631337) was modified by inserting EF1a-TagRFP-2A-tet3G with tetON-NKX2-1 CDS, or by inserting EF1a-mNeonGreen-2A-tet3G with tetON-TFAP2C CDS. For generating NKX2.1 variants, naturally occurring mutations in NKX2.1 binding domain region was selected from Leiden Open Variation Database 3.0^14^ (www.lovd.nl/3.0) and two previously reported clinical cases^12,13^ – 1 amino acid deletion^14^ (p.R162del), two nonsense point mutations^14^ (p.Q175* and p.R178*), and two missense point mutations^12,13^ (p.I207F and p.I207M). NKX2.1 CDS harbouring each mutation was amplified and inserted by Infusion (638909, Takara) cloning into the tetON-NKX2.1/EF1a-TagRFP-2A-tet3G Dox-inducible lentiviral vector. NKX2.1 CDS lacking the entire DNA binding domain was inserted into the EF1a-TagRFP-2A-tet3G Dox-inducible lentiviral vector by Infusion cloning.

For the NKX2.1 knock-down experiment, a modified Dox-inducible CRISPRi vector was gifted^23^; N-terminal KRAB-dCas9 (a gift from Bruce Conklin, Addgene plasmid # 73498) fused with a destabilising domain, dihydrofolate reductase (DHFR) sequence that is only stabilised by trimethoprim (TMP) treatment, was sub-cloned into the EF1a-TagRFP-2A-tet3G Dox-inducible lentiviral vector^23^. Treatment of 2 µg/ml Dox (Merck, D9891) with 10 nmol/L TMP (Merck, 92131) in the SN medium stabilize the functional KRAB-dCas9 protein. Three gRNAs targeting NKX2.1^24^ were individually subcloned into gRNA lentivirus as follows: gRNA-1; 5’-GTCTGACGGCGGCAGAAGAG-3’, gRNA-2; 5’-GGACCAACAGTGCGGCCCCA-3’, gRNA-3; 5’-GAAATGAGCGAGCGAGTCTG-3’. Single cells dissociated from organoids were infected and the infected cells were sorted by TagRFP and/or mNeonGreen fluorescent signal using FACS (Fig. 5) after 48 h of infection. The sorted TagRFP^+^ and/or mNeonGreen^+^ cells were cultured in the Matrigel in the absence of Dox or TMP for 1 week. After the cells were grown into a typical organoid, the Dox and/or TMP were added and culture continued for additional 2 weeks.

### Immunostaining of organoids and lung tissues

For immunostaining of human lung tissue sections, the lungs were fixed in 4% paraformaldedyde (PFA; Merck, 158127) overnight, washed in PBS and 15%, 20% and 30% sucrose (w/v) in PBS before embedding in Optimum Cutting Temperature (OCT) medium (Merck, F4680). 12 μm thick frozen sections were collected and permeabilised using 0.3% Triton-X in PBS for 15 min. Antigen retrieval was performed by heating the slides in 10 mM Na-Citrate buffer at pH 6.0 in a microwave for 5 min. Then slides were treated with blocking solution containing 5% NDS, 1% Bovine Serum Albumin (BSA), 0.1% Triton-X in PBS at room temperature for 1 h.

For whole-mount immunostaining of lung organoids, the Matrigel was completely removed from the cultured organoids using Cell Recovery Solution (Corning, 354253) and fixed in 4% PFA for 30 min on ice. After rinsing in PBS washing solution containing 0.2% (v/v) Triton X-100 and 0.5% (w/v) BSA, the samples were transferred to a round-bottom 96 well plate and incubated in permeabilization/blocking solution containing 0.2% (v/v) Triton X-100, 1% (w/v) BSA, and 5% normal donkey serum (NDS) in PBS, overnight at 4°C.

For primary antibody treatment, the following antibodies were treated to the organoids and the tissue slices at 4°C overnight: proSFTPC (1:200; Merck, AB3786), E-cadherin (1: 500; Thermo Fisher Scientific, 13-1900), NKX2.1 (1:200; Merck, 07-601), TFAP2C (1:200; Abcam, ab218107), CD44 (1:200; Thermo Fisher Scientific, 17-0441-82), CD36 (1:200; Proteintech, 18836-1-AP), alpha-smooth muscle actin (1:500; Thermo Fisher Scientific, MA1-06110), ACE2 (1:100; Abcam, ab108252), AXIN2 (1:200; R&D Systems, MAB6078), PDPN (1:200; R&D Systems, AF3670), CD31 (1:200; Abcam, ab9498), PDGFRA (1:200; Cell Signaling Technology, 3174), TP63 (1:200; Cell Signaling Technology, 13109), SOX2 (1: 500, Bio-techne, AF2018), SOX9 (1: 500, Merck, AB5535), LAMP3 (1:100; Atlas Antibodies, HPA051467), HTII-280 (1:200; Terracebiotech, TB-27AHT2-280), CD141 (1:100; PE-conjugated; BioLegend, 344104), ZO-1 (1:200; Invitrogen, 40-2200) and KI67 (1:200; BD Biosciences, 550609). After three washes with PBS, 97% (v/v) 2’−2’-thio-diethanol (TDE, Sigma, 166782) was treated for clearing. Images were collected under Leica SP8 confocal microscope.

### *In situ* hybridization chain reaction (*in situ* HCR)

*In situ* HCR v3.0 was performed according to the manufacturer’s procedure (Molecular Instruments^25^). Probes were designed according to the protocol and amplifiers with buffers were purchased from Molecular Technologies. Sequence information of the probes for detecting *SFTPC, WNT2, NOTUM, AXIN2, SOX9, FGFR4* and *TFAP2C* mRNA targets is listed in Supplementary Table 3. Briefly, frozen human tissue sections were cut at 20 μm from lungs fixed overnight in 4% PFA in DEPC-treated PBS and processed to cryoblocks. Lung sections were carefully rinsed in nuclease-free water, followed by 10 μg/mL proteinase K treatment (Thermo Fisher Scientific, AM2546), and 2 pmol of each probe was treated at 37°C overnight. After washing, the tissue was incubated with 6 pmol of the amplifiers at room temperature overnight for amplification. The amplifiers, consisting of a pair of hairpins conjugated to fluorophores, Alexa 546, 647, or 488, were snap-cooled separately and added at final 0.03 µM to the tissue. After removing excess hairpins in 5X SSC (sodium chloride sodium citrate) buffer containing 0.1% Triton X-100, nuclei were counter-stained with DAPI.

To combine *in situ* HCR with antibody immunostaining, the frozen human tissue sections from 20 μm up to 100 μm thickness were permeabilised using 0.3% Triton-X in DEPC-treated PBS for 20 min at room temperature. Then the tissues were treated with blocking solution containing 5% NDS, 1% BSA, 0.1% Triton-X in DEPC-treated PBS at 4°C for 3 h. After rinsing with cold DEPC-treated PBS, treated with a primary antibody against ACTA2 (1:500; Thermo Fisher Scientific, MA1-06110) for 24 h, followed by a secondary antibody treatment (1:500; Thermo Fisher Scientific, A10036) at 4°C overnight. After the tissue was washed three times in the DEPC-treated PBS at room temperature, 2 pmol of each *in situ* HCR probes for targeting *SFTPC* and *NOTUM* was treated at 37°C overnight without 10 μg/mL proteinase K treatment to preserve the antibody immunofluorescence. After washing, the tissues were incubated with 6 pmol of the amplifiers at room temperature overnight. Following three times of rinsing in in 5X SSC buffer containing 0.1% Triton X-100, nuclei were stained with DAPI. Finally, the tissues were processed to 2’-2’-thio-diethanol (TDE, Sigma, 166782) for clearing and mounting: 10 %, 25 %, 50 % (v/v) TDE in 1x DEPC-treated PBS for 1 hr and 97% TDE overnight at 4°C. Images were collected under Leica SP8 confocal microscope.

### RNA extraction, cDNA synthesis, qRT-PCR analysis, and bulk RNA-seq

Organoids were removed from the Matrigel and lysed. Total RNA was extracted according to the RNeasy Mini Kit (Qiagen, 74004) protocol. For cells freshly purifed from human lung tissues were directly lysed using 100 μl lysis buffer from PicoPure™ RNA Isolation Kit (Thermo Fisher Scientific, KIT0204). First Strand cDNA synthesis was performed using High-Capacity cDNA Reverse Transcription Kit (Applied Biosystems, 4368814). Then, cDNA was diluted 1:50 for qRT-PCR reaction (SYBR Green PCR Master Mix; Applied Biosystems, 4309155). Primer sequence information is listed in Supplementary Table 4. Data is presented as fold change, calculated by ddCt method, using *ACTB* as housekeeping reference gene. For bulk RNA-seq, RNA quality was validated on Agilent 2200 Tapestation. The RNA-seq libraries were generated at the Cancer Research UK Cambridge Institute and sequenced on an Illumina HiSeq 4000. A list of differentially expressed genes was extracted using the counted reads and R package edgeR^26^ version 3.16.5 for the 3 pairwise comparisons (Supplementary Table 1). GO biological processes term enrichment, KEGG pathway, and gene set enrichment analysis were performed using DAVID^27^, Enrichr^28^, and R package fgsea package^29^, respectively.

### Immunoblotting

The organoid samples were harvested and lysed (RIPA buffer; Merck, R0278) after complete removal of the Matrigel and run on 12.5 ∼ 20 % SDS PAGE gels. Proteins on the gels were transferred onto PVDF membrane using BioRad Mini Trans-Blot system (BioRad, Mini Trans-Blot^®^ Cell). The membranes were washed with pure water and blocked with 5% skimmed milk in 0.1% Tween-20/PBS (PBST) for 30 min at room temperature. Membranes were incubated with primary antibodies against NKX2.1 (1:200; Merck, 07-601), proSFTPC (1:1000; Merck, AB3786), mature SFTPC (1:1000; Seven Hills Rioreagents, WRAB-76694), mature SFTPB (1:1000; Seven Hills Rioreagents, WRAB-48604), and GAPDH (1:5000; Abcam, ab8245) in the blocking buffer overnight at 4 °C. After washing with PBST, secondary antibodies conjugated with fluorescence dyes (1:5000; anti-mouse IRDye® 800CW and anti-rabbit IRDye® 680RD; Abcam, ab216774 and ab216779, respectively) were treated at room temperature for 3 h. The membranes were washed in PBST and developed using Li-Cor Odyssey imaging system.

### Chromatin immunoprecipitation

Chromatin immunoprecipitation (ChIP) was performed according to SimpleChIP® Chromatin immunoprecipitation protocol (Cell Signaling Technology, 9002). In brief, the organoids were harvested and enzymatically dissociated into single cells using TrypLE Express Enzyme (Thermo Fisher Scientific, 12605010). Then the cells were crosslinked with 1% formaldehyde for 15 min at room temperature and the reaction was quenched by glycine at a final concentration of 0.125 M. Chromatin was digested with 1 μl MNase (Cell Signaling Technologies, 10011S) for 20 min at 37°C, followed by sonication for 12 cycles of 30 seconds on and 30 seconds off using Biorupter (Diagenode, UCD-300), to length of an average size of 150-900 bp. 5 μg of digested chromatin samples was treated with antibodies against rabbit IgG (1:100; Cell Signaling Technology, 2729) or NKX2.1 (1:100; Merck, 07-601). The amount of immunoprecipitated DNA was quantified by qPCR using primers specific for promoter regions of SFTPC, LAMP3, and SLC34A2. Fold enrichment values are presented as the fold-change over the level of ChIP with negative control IgG antibody (ChIP signal/IgG signal). Sequence information of the primers for targeting SFTPC, LAMP3, and SLC34A2 promoter regions is listed in Supplementary Table 4.

### Bulk ATAC-seq

Genome-wide chromatin accessibility of lung organoids was assessed as previously described^30^. In brief, 50,000 cells were harvested from organoids and lysed in lysis buffer (10 mM Tris-HCl, pH 7.4, 10 mM NaCl, 3 mM MgCl2, 0.1% (v/v) IGEPAL CA-630). The lysate was treated in 50 μL reactions with Nextera TDE1 transposase (Illumina, 15027865) for 30 min at 37°C. The purified DNA was amplified and indexed using Nextra DNA CD Indexes (Illumina, 20018707), and size distribution of the DNA libraries was analysed using High-sensitivity Qubit dsDNA Assay Kit (ThermoFisher, Q32851) and Agilent 2200 Tapestation. The libraries were sequenced on an Illumina HiSeq 4000. Peak calling was done using MACS2 algorithm^31^ (version 2.1.1) and further processed to extract differential peaks (Supplementary Table 2). Then, the differential peak data was further used for analysing transcription factor motifs using HOMER^32^ software in combined with RNA-seq data.

### Electron microscopy imaging

The organoid samples were fixed in 2 % formaldehyde/2 % glutaraldehyde in 0.05 M sodium cacodylate buffer (NaCAC), pH 7.4, containing 2 mM calcium chloride (Merck, C27902) overnight at 4°C. After washing in 0.05 M NaCAC at pH 7.4, the samples were osmicated for 3 days at 4°C. After washing in deionised water (DIW), the samples were treated twice with 0.1 % (w/v) thiocarbohydrazide (Merck, 223220) in DIW for each 20 min and 1 h at room temperature in the dark, followed by block-staining with uranyl acetate (2 % uranyl acetate in 0.05 M maleate buffer pH 5.5) for 3 days at 4°C. Then, the samples were dehydrated in a graded series of ethanol (50%/70%/95%/100%/100% dry) 100% dry acetone and 100% dry acetonitrile, three times in each for at least 5 min. Next, the samples were infiltrated with a 50:50 mixture of 100% dry acetonitrile/Quetol resin (TAAB, Q005) without BDMA (TAAB, B008) overnight, followed by 3 days in 100% Quetol without BDMA. The sample was infiltrated for 5 days in 100% Quetol resin with BDMA, exchanging the resin each day. The Quetol resin mixture is: 12 g Quetol 651, 15.7 g NSA (TAAB, N020), 5.7 g MNA (TAAB, M012) and 0.5 g BDMA. Samples were placed in embedding moulds and cured at 60°C for 3 days.

Thin sections were cut using an Ultracut E ultramicrotome (Leica) and mounted on melinex plastic coverslips. The coverslips were mounted on aluminium SEM stubs using conductive carbon tabs and the edges of the slides were painted with conductive silver paint. Then, the samples were sputter coated with 30 nm carbon using a Quorum Q150 T E carbon coater and imaged in a Verios 460 scanning electron microscope (FEI, Thermo Fisher Scientific) at 4 keV accelerating voltage and 0.2 nA probe current in backscatter mode using the concentric backscatter detector in immersion mode at a working distance of 3.5-4 mm; 1,536 x 1,024 pixel resolution, 3 μs dwell time, 4 line integrations. Stitched maps were acquired using FEI MAPS software using the default stitching profile and 10% image overlap.

## Data availability

All bulk RNA-seq and ATAC-seq data generated have been deposited in NCBI’s Gene Expression Omnibus and are accessible through GEO Series accession number GSE178529.

### Statistical analysis

Data are expressed as average ± standard deviation (SD). Statistical significance was evaluated by unpaired student’s *t* test, 1- or 2-way ANOVA with Tukey/Bonferroni/ Dunnett comparison multiple comparison post-test; ns: not significant, \**P*<0.05, \*\**P*<0.01, \*\*\**P*<0.001, and \*\*\*\**P*<0.0001.

